# Mechanical control of tissue growth during limb regeneration

**DOI:** 10.1101/2025.04.07.647008

**Authors:** Sandra Edwards-Jorquera, Alberto Ceccarelli, Raimund Schlüßler, Nivedha Murali Shankar, Angela Muñoz Ovalle, Kyle Gentleman, Uwe Freudenberg, Carsten Werner, Osvaldo Chara, Anna Taubenberger, Tatiana Sandoval-Guzmán

## Abstract

The axolotl is a highly regenerative species, capable of restoring full limbs, regardless of the amputation site. However, the regeneration rate is adjusted with the plane of amputation along the proximo-distal (PD) axis, leading to equivalent regeneration times regardless of the extent of tissue removal. We hypothesized that this phenomenon could be partly explained by differences in tissue mechanical properties. In this work, we describe tissue growth mathematically and evaluate cell cycle parameters of regenerating limbs amputated at different levels along the PD axis, demonstrating a linear correlation between the cell cycle length and the amputation site during early regeneration phases. We show as well, that blastema cells require their endogenous context to retain such proliferation differences. We measured mechanical properties in regenerating limbs with *in vivo* optical and standard indentation-based techniques and demonstrated that distal blastema cells are stiffer than proximal ones. Accordingly, we demonstrated that axolotl cells decrease their proliferation with increased extracellular matrix stiffness *in vitro*. Next, we evaluated the activity of the mechanotransducers YAP/TAZ *in vivo* by using a *GTIIC*-based reporter line combined with target gene expression data, which indicated that their activity peaks during the blastema stage, with higher activity after proximal amputations. Hence, our findings strongly suggest a mechanical dependence for the position-dependent regulation of cell proliferation during axolotl limb regeneration, where YAP/TAZ likely plays a role in the mechanotransduction mechanism.

## INTRODUCTION

In multicellular organisms, tissue growth is dynamically yet precisely regulated throughout life. During development, processes such as cell proliferation, differentiation, and migration are systemically activated and coordinated to ensure the formation of appropriately sized and functional organs. In most adult organs, size is maintained by a slow and steady cell turnover. However, upon tissue loss, cell proliferation can be reactivated near the injury site, leading to localized growth rates that are independent of organismal development, thus allowing the regeneration of lost body parts (reviewed in ^1^).

A great model species to study growth regulation in tetrapods is salamanders, as they can regrow full limbs following amputation, and the regenerating limb grows until it catches up with the development of the intact contralateral one ^2^. Limb regeneration requires the formation of the blastema, a structure containing precursor cells for the formation of lost tissues, which is covered by a specialized epithelium called the apical epithelial cap (AEC). Through overlapping phases of growth, cell differentiation and pattern formation, the blastema rebuilds the missing limb (reviewed in ^3^). Amputation anywhere along the salamander limb yields regeneration of only the missing portion, which is distal to the level of amputation. This indicates that mature cells within the limb stump retain a memory of their location along the proximo-distal (PD) axis, which blastema cells interpret to regenerate only the missing parts – a property termed positional identity ^4^. Strikingly, in salamanders, following the amputation of nearly the entire limb on one side and only a digit on the contralateral, regeneration of both structures is completed in the same time, independent of the tissue volume to be reformed ^5^. Thus, the growth rate depends on the amputation level. Similar observations were reported for other appendages in salamanders ^6–8^, as well as different body parts across species, including frogs, teleosts, lizards, and starfish ^9–13^. This interesting phenomenon raises the question of how does the amputation site translate into controlled growth?

Zebrafish sense the amputation position through a gradient of tissue mechanical tensions coupled to a travelling density wave in the caudal fin basal epithelium, which is associated to position-dependent levels of reactive oxygen species (ROS) ^14^. Moreover, it is proposed that this initial mechanical wave and brief peak in ROS are linked to different expression levels of Fibroblast Growth Factor (FGF) in epidermal cells ^10^ via ERK signaling in osteoblasts ^15^. The amputation-dependent growth phenomenon in regenerating zebrafish caudal fins has also been related to a regeneration-specific H^+^ efflux ^16^, lysosomal acidification, and mTORC1 activity ^17^. In *Xenopus* embryos, it was shown that inhibiting the transcription factor TEAD4 decreases growth differences between anteriorly and posteriorly amputated tails ^13^. It is yet to be determined whether there is an interplay between these mechanisms, and whether they are conserved in tetrapod limbs or are tissue- and species-specific.

In the axolotl, it has already been established that limb positional information is determined in connective tissue cells ^18,19^ through segment-specific epigenetic modifications in *Meis* and *Hoxa13* gene *loci*, which are accompanied by the basal expression of some positional genes^20^. It has also been proposed that positional identity is encoded at the cell surface during regeneration, as engulfment assays ^21–23^ and Steinberg’s differential adhesion hypothesis ^24^ suggest that distal blastema cells have stronger intercellular adhesions than proximal ones. Finally, extracellular matrix (ECM) components have been reported to be distributed across a gradient within blastemas and, therefore, could contribute to positional information as well ^25^.

Tissue surface tension depends on the ratio of adhesion to cortical tension of individual cells ^26,27^, as well as the characteristics of the ECM ^28^. These factors are responsible for force transmission to and between cells, thereby controlling signaling pathways that regulate stem cell self-renewal and differentiation ^29,30^. Accordingly, mechanical properties drive processes such as cell proliferation, differentiation, and migration in multiple contexts ^29,31–34^, and have been shown to greatly influence tissue growth and morphogenesis during development ^31,33–35^, loss of stemness during aging ^32^, and the progression of pathologies like fibrosis and cancer ^36,37^. More recently, it has been implied that physical forces play relevant roles in stem cell function during regeneration as well ^14,30,38–40^. Therefore, the differences in cell adhesion and ECM along the PD axis may be associated with positionally regulated tissue mechanical properties in regenerating axolotl limbs. Whether such mechanical differences exist is, thus far, unknown. Furthermore, the impact of mechanical properties on the differential growth rates observed during regeneration between proximally and distally amputated limbs has not been explored yet.

The best described pathway capable of integrating a broad spectrum of cues, including mechanical, and transducing them into the controlled growth, is the Hippo signalling pathway, which has thus been designated as a central growth control mechanism in multicellular organisms (reviewed in ^1^). This pathway culminates in the phosphorylation of the transcriptional coactivators YAP and TAZ, leading to their cytoplasm retention or proteasomal degradation. When not inhibited, these proteins translocate to the nucleus, where they bind to TEAD transcription factors, inducing the expression of a wide range of genes associated with cell proliferation, differentiation, and survival – for simplicity, we will hereafter treat them as equivalent proteins, even though both distinct and overlapping functions of YAP and TAZ have been described ^41^. Accordingly, restricting YAP activity has been shown to limit growth during regeneration in several species ^13,42–44^.

Here, we studied the implication of mechanical properties in growth regulation along the limb PD axis during regeneration in the axolotl. We mathematically modelled tissue growth, revealing a linear correlation with the amputation plane during early phases of regeneration, and we demonstrate that differences in proliferation between proximal and distal blastema cells are non-cell autonomous. We describe mechanical properties with optical and standard indentation-based techniques and show that distal blastemas are stiffer than proximal ones. Furthermore, our results indicate that cell proliferation differences along the limb PD axis are correlated with YAP/TAZ activity and opposed by tissue stiffness levels and cell differentiation times, suggesting that growth rate is mechanically regulated during axolotl limb regeneration in a process mediated by YAP/TAZ. Thus, our work illustrates how a position-related growth program can be regulated by mechanical forces during regeneration.

## RESULTS

### Limb growth rate is directly correlated with the amputation plane

The evidence for tissue growth rate depending on the amputation level along the limb PD axis ^5–7^ has been mostly descriptive and only quantified via length measurements in the newt *Notophthalmus viridescens* ^7^. Here, we measured the limb’s 2-dimensional dorsal projection to quantify limb size in the axolotl (*Ambystoma mexicanum*). Moreover, we coupled our measurements to a mathematical model to calculate the growth rate and mean cell cycle length. For this, we amputated limbs at the upper arm, lower arm, or digit and followed regeneration progression over time in two groups of differently sized animals (Figures 1A, S1A). To obtain a reliable assessment of tissue growth, we measured limb area from the joint proximal to the amputation site, of both amputated and contralateral intact limbs, and showed that, as expected, proximally amputated limbs grow faster than those distally amputated (Figures 1B), irrespective of animal size (Figures S1B). We also found that the relative limb size – the area of the regenerating limb with respect to the intact contralateral one – increases at a rate that is independent of the amputation plane (Figures 1C, S1C). Weekly growth calculations indicate that proximally amputated limbs grow larger tissue volumes *per* week than distally amputated ones (Figure 1D).

**Figure 1:**
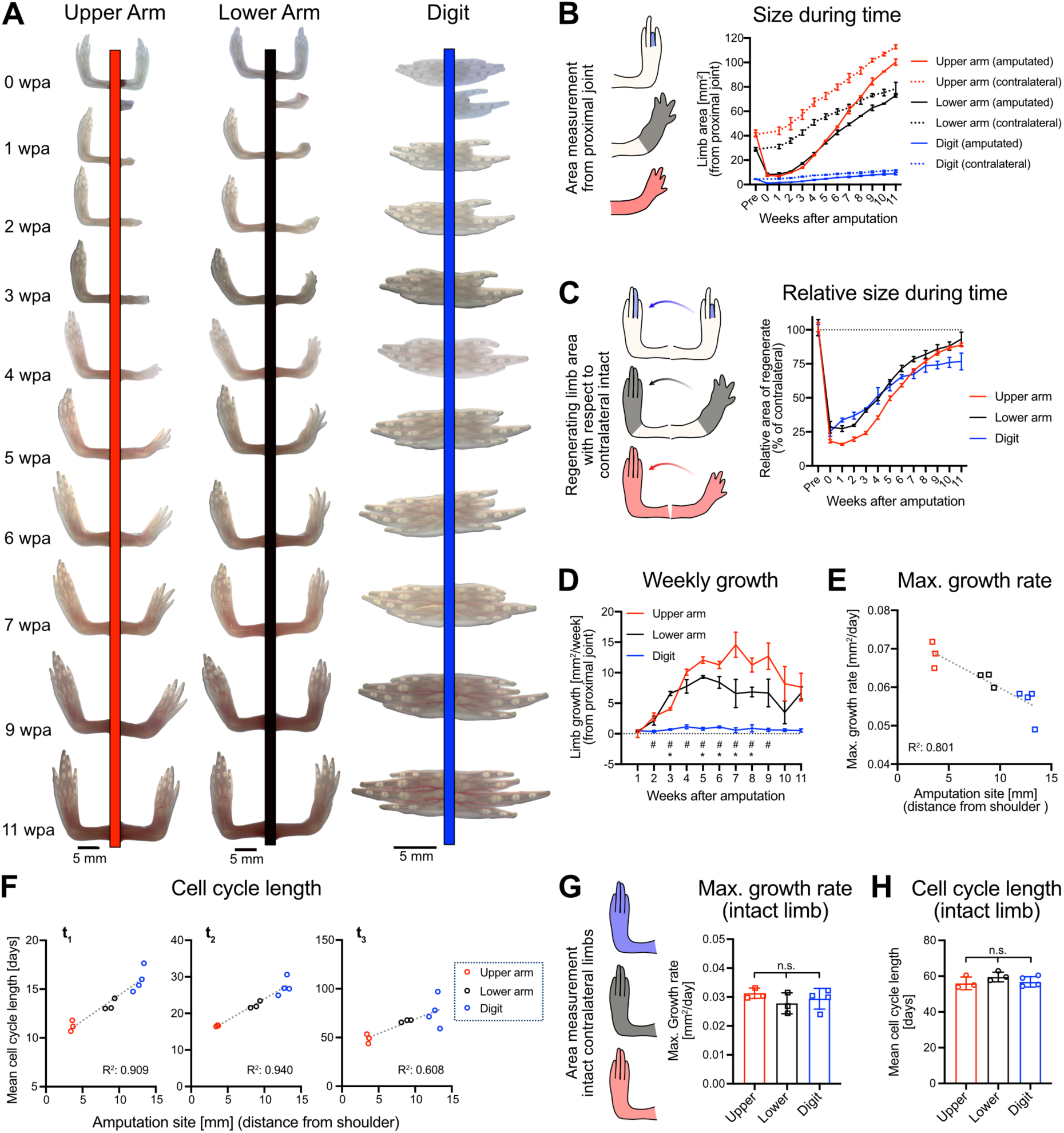
Tissue growth during regeneration is faster after proximal amputations. 4-month-old animals were amputated at the upper arm, lower arm or digit and regeneration was assessed weekly. **A)** Representative images of amputated (right) and intact (left) limbs during 11 weeks. wpa: weeks post-amputation. **B)** Left: Tissue area was measured from the joint most proximal to the amputation site. Right: Limb area from regenerating and intact limbs (n ≥ 3 animals/condition). **C)** Relative area of regenerating limb with respect to intact contralateral. **D)** Weekly growth calculated from values in *B*. Two-way ANOVA with Tukey’s multiple comparisons test, * *p* < 0.05 (Upper arm *vs.* digit), ^#^ *p* < 0.05 (Upper arm *vs.* Lower arm). **E)** Correlation of maximum growth rate with respect to amputation site. **F)** Mean cell cycle duration with respect to amputation site during different phases of regeneration. The regeneration time course was divided into 3 equivalent segments (t_1_, t_2_ & t_3_). **G-H)** Maximum growth rate (*G*) and mean cell cycle length (*H*) of intact contralateral limbs (measured from shoulder). One-way ANOVA with Kruskal-Wallis multiple comparisons test, n.s. no statistically significant differences. For *B-D, G-H:* Mean ± SD is shown. For *E-H*: Each dot represents one animal. For *E-F*: Lines represent linear regression and coefficient of determination (R^2^) is indicated.

We mathematically described the dorsal projection growth with a logistic regression model, which fitted well to our experimental data (Figures S2A, S2C). This model revealed that the maximum growth rate has an inverse linear correlation with the amputation site, expressed as distance from the shoulder (Figure 1E). However, in small animals, this correlation excludes the digits (Figure S1D). We hypothesize that digits may still be growing at a higher rate in small animals due to the recent culmination of limb development prior to amputations ^45^. Throughout the regeneration time course, the mean cell cycle length is shorter after proximal amputations when compared to distal ones (Figures S2B, D). However, the mean cell cycle length changes notably during the process, becoming increasingly larger towards the end of it (Figures S2B, D), which was expected, as the increase in limb size during regeneration is not linear (Figure 1B, S1B). Therefore, we subdivided the process into 3 equivalent phases (Figure S2E) and determined the mean cell cycle length during each of them. This subdivision shows that there is a linear correlation between the amputation site and the cell cycle length during early and intermediate regeneration stages, which recedes at the later stage (Figure 1F, S1E).

As a proxy to determine whether amputations affected organismal growth, we compared the contralateral intact limbs of the axolotls amputated at different levels, as well as intact siblings. Akin to the amputated limb calculations, we determined the maximal growth rate (Figures 1G, S1F) and the mean cell cycle length (Figures 1H, S1G) between conditions, which indicate that all amputated animals grow equivalently to unamputated ones. These values strongly suggest that regeneration does not delay body growth and development in the axolotl, contrary to observations in insects ^46,47^. Additionally, to determine whether distinct limb segments grow allometrically, we compared intact limb growth from different levels along the PD axis (Figure S2F-G). We found differential growth by animal size, with larger animals having a higher maximum growth rate in lower arm segments, which is not observed in younger animals (Figure S2F). In addition, in larger animals, no differences in cell cycle length were detected between limb segments, whereas in smaller animals, it is longer in the digits (Figure S2G).

Altogether, we demonstrate that limb growth differences along the PD axis commence at early regeneration stages and are directly correlated with the amputation site, with proximally amputated limbs growing faster and presenting shorter mean cell cycle lengths than distally amputated ones.

### Cell proliferation and differentiation during regeneration are inversely regulated along the PD axis during regeneration

To confirm the model’s estimations, we used animals expressing a fluorescent cell cycle reporter (*AxFUCCI*) ^48^ to evaluate cell proliferation at different regeneration stages: AEC, blastema, palette, and digit patterning (Figure 2A). We quantified the proportion of proliferating cells, evidenced by mCherry expression in wholemount. Our results indicate that blastemas derived from proximal amputations (hereafter referred to as proximal blastemas) contain a significantly higher proportion of proliferating cells in the mesenchyme, and a similar trend was observed in the epithelium at later stages (Figure 2B). These results suggest that the shorter mean cell cycle length during early phases of regeneration in proximally amputated limbs (Figure 1F, S1E) may be explained, at least in part, by their higher proportion of proliferating cells. Complementarily, we evaluated EdU incorporation in tissue sections from adult animals (Figure 2C). Consistent with *AxFUCCI* measurements, the EdU incorporation assessment indicates a higher proportion or proliferating cells in proximal blastemas, both in the mesenchyme as well as the epithelium (Figure 2D). Furthermore, we used the *Brainbow2.1* transgenic line ^49^ to induce the random expression of four different fluorescent proteins (Figure S3A). Thus, by assuming that each cell acquired a unique color combination prior to amputation, we used individual clone sizes at different regeneration time points as an indication of cell proliferation rate at different regeneration time points (Figure S3B). With this system, we show that proximal blastemas have larger clone sizes than distal ones, suggesting that cells have undergone more rounds of duplication. Such differences are accentuated at later regeneration stages (Figure S3C).

**Figure 2:**
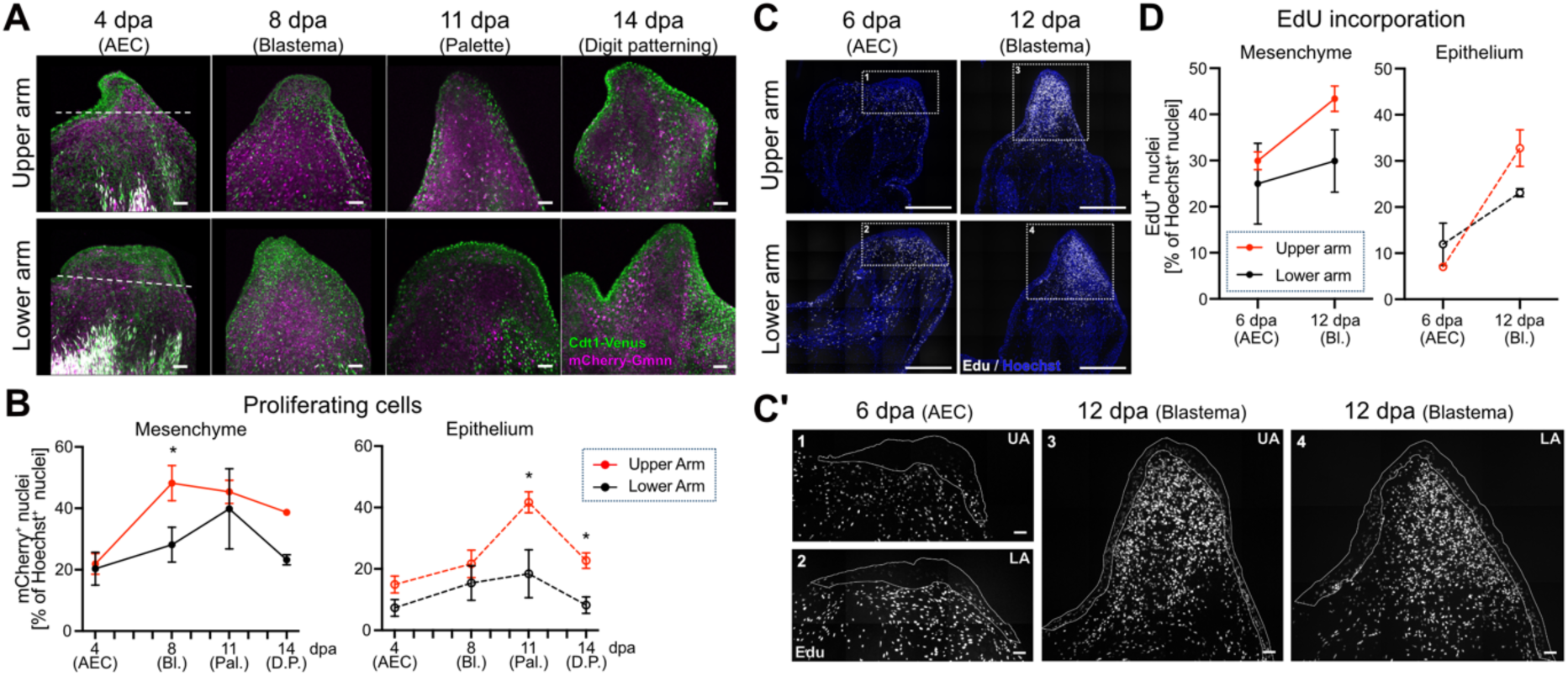
Proximal blastemas have a higher proportion of proliferating cells than distal ones. Animals were amputated at the upper and lower arm levels, after which limbs were collected at different regeneration timepoints. **A)** Representative images of regenerating limbs of *AxFUCCI* animals ^48^ after tissue clearing. Merged signal of Cdt1-Venus (green) and mCherry-Gmnn (magenta) are shown. Scale bar: 100 µm. **B)** Proportion of proliferating mCherry^+^ cells in the mesenchyme and epithelium. Mean ± SD is shown (n = 3 animals/condition). Two-way ANOVA, Sidak’s multiple comparisons test, * *p* < 0.05. **C)** Animals were injected with EdU 4h prior to tissue collection. Representative images of tissue sections with labelled EdU+ (white) and all nuclei (Hoechst, blue). Scale bar: 1 mm. **C’**) Enlarged sections from dotted rectangles in *C* showing EdU signal and outlined epithelium. Scale bar: 100 µm. **D)** Proportion of EdU^+^ nuclei in the mesenchyme and epithelium. An average of 8 tissue slices *per* animals we used (n = 2 animals/condition). Mean ± SD is shown. dpa: days post amputation.

Using these different but complementary approaches, we show that growth differences already become apparent at the blastema stage, much earlier than previously reported ^7^. Interestingly, we also detected differences in digit patterning dynamics, which occurred earlier following distal amputations (Figure 1A, 3-4 weeks post-amputation (wpa) and Figure 2A, 14 days post-amputation (dpa)). These observations, together with our proliferation assessment (Figure 2 and S3), suggest that there may be differences in the regulation of cell cycle exit and differentiation in blastemas along the PD axis. To address this hypothesis, we histologically analyzed limbs amputated at the upper and lower arm at different stages of regeneration by staining the connective tissue with Movat’s pentachrome (Figure 3A-F). We detected that, at the AEC stage, the epithelial layer is thicker – a sign of AEC maturation – and located in the center of the limb after an upper arm amputation, compared to the lower arm amputation where it is thinner but broader (Figure 3A-B). This difference can also be observed at an equivalent stage in younger animals imaged in wholemount (Figure 2B, 4 dpa). Considering that nerve signals are crucial for AEC establishment ^50^ and that nerve fibers diminish along the PD axis, we hypothesized that a proximal amputation will result in more nerve signals than distal ones. Using a transgenic *βIII-Tubulin* reporter line to label neurons with mCherry ^51^, we imaged regenerating limbs in wholemount and detected that indeed nerve bundles were thicker and more concentrated in the central part of the arm (around the humerus) in proximally amputated limbs, as compared to distally amputated ones, where nerve terminals spread out more (around the radius and ulna) (Figure S4).

**Figure 3:**
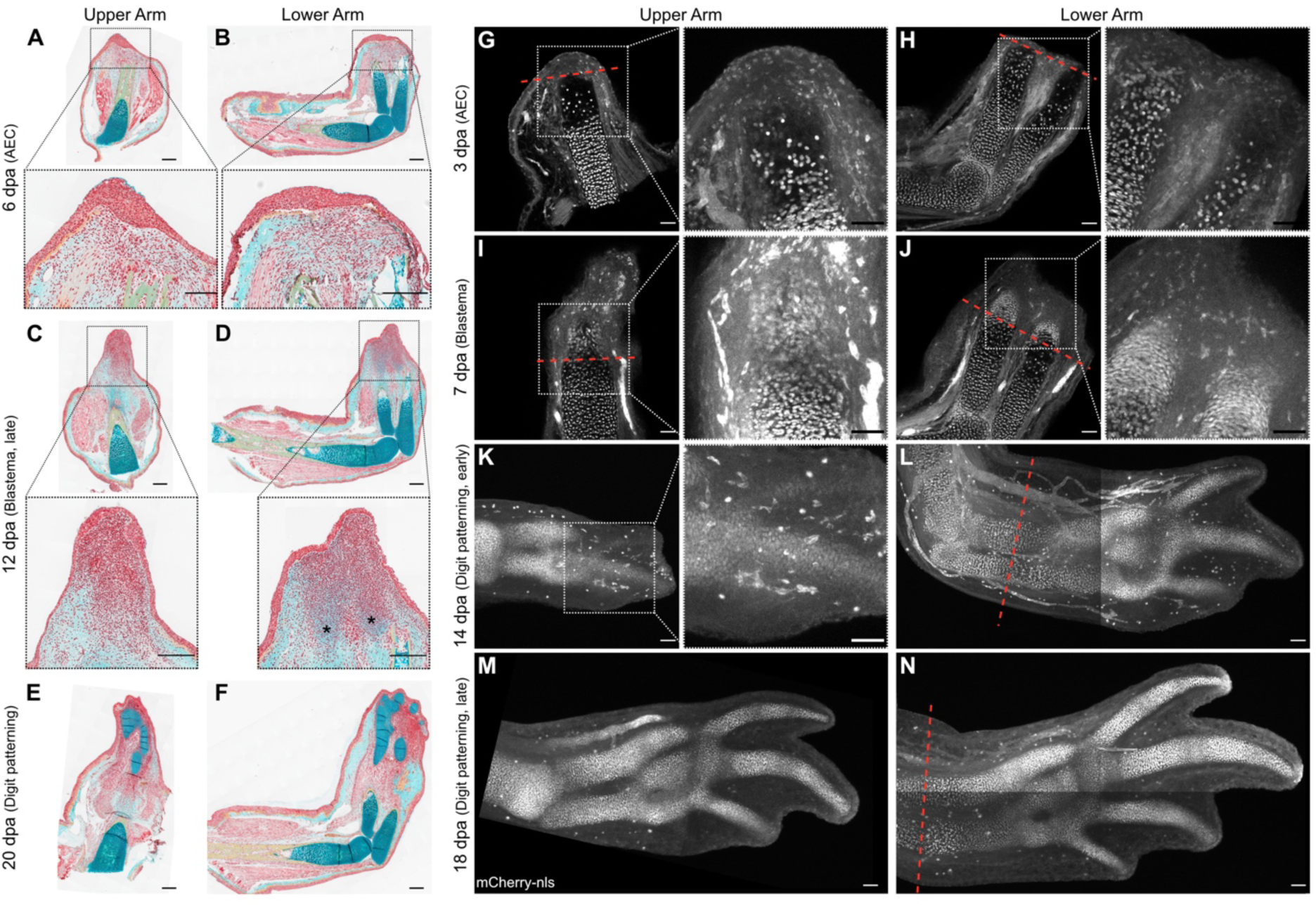
Tissue morphogenesis during regeneration occurs faster after distal amputations. Limbs were amputated at the upper and lower arm levels, after which they were collected at different regeneration timepoints. **A-F)** Movat’s pentachrome stained tissue sections: red (nuclei, muscle), light blue (connective tissue), blue/green (cartilage), yellow (mineralized tissue). Scale bar: 1 mm. **A-B)** Apical epithelial cap (AEC) revealing differences in epithelial layer between upper and lower arm amputations. **C-D)** Late blastema stage indicating the faster morphogenetic process occurring after distal amputations, including early cartilage condensation (indicated by *). **E-F)** Digit patterning stages showing the difference in hand width and digit number between amputation planes. **G-N)** Wholemount fluorescent imaging of optically cleared limbs from *Sox9:Sox9-T2A-mCherry* expressing animals revealing skeletal elements. Dashed red line indicates the amputation plane. Scale bar: 100 µm. For *G-J*: Overview: one optical plane is shown, Close up: z-stack enclosing entire thickness of the bone. For *K-N*: z-stack of entire limbs. Dpa: days post-amputation.

At the late blastema stage, we can observe in the lower arm a group of cells with flattened nuclei organized in columns, akin to early stages of cartilage condensation ^52^, which are not visible in the upper arm (Figure 3C-D). To study cartilage cell differentiation, we evaluated the distribution of SOX9-expressing cells (Figure 3 G-N), as this transcription factor is among the first tissue-specific genes to be expressed during limb regeneration ^18^, making it a good marker for early cell differentiation. Therefore, we used a *Sox9*-transgenic knock-in line, where SOX9-expressing cells are labelled with nuclear mCherry ^52^, and evaluated the distribution of mCherry^+^ nuclei during different stages of limb regeneration in wholemount (Figure 3G-N). As expected, we did not detect any condensation of SOX9-expressing cartilage precursor cells at the AEC stage (Figure 3G-H), but in the blastema, SOX9-expressing cells begin to condense distally to the bone. However, following a lower arm amputation, these cells display a columnar morphology and organization (Figure 3I-J), an indication of a later stage of cartilage differentiation ^53^, which is not observed in the upper arm blastema. At the digit patterning phase, the differences in cartilage differentiation are accentuated, revealed by both cartilage staining in tissue sections (Figure 3E-F) and SOX9-expressing cells in wholemount (Figure 3K-N).

Taken together, proximal blastemas have a higher proportion of proliferating cells and delayed differentiation compared to distal blastemas. These observations strongly suggest that cells in distal blastemas are signaled to differentiate and exit the cell cycle sooner, thus reducing the pool of remaining proliferating cells.

### Blastema mechanical properties are position-dependent

Changes in stem cell self-renewal and differentiation can be modulated by the mechanical environment ^29,30^. Therefore, we hypothesized that the differences detected in cell proliferation and differentiation in limb blastemas along the PD axis are associated with different mechanical properties. To test this, we mechanically probed axolotl limbs at different stages of regeneration.

We characterized the tissue mechanical properties by using two different but complementary techniques: atomic force microscopy (AFM) ^54^ and Brillouin confocal microscopy ^55^, both of which were recently optimized for the measurement of regenerating axolotl limbs ^40,56^.

To mechanically probe internal tissues with AFM, we sectioned regenerating limbs with a vibratome (Figure 4A) and performed multiple indentation tests using a cell-sized spherical indenter on different regions within the epithelium and the mesenchyme. In general, we found that epithelial regions have significantly higher apparent Young’s moduli than the mesenchyme. Moreover, distal blastemas have higher apparent Young’s moduli than proximal ones in both tissue types (Figure 4B), indicating increased tissue stiffness. The differences observed between proximally and distally amputated limbs at the blastema stage are not present during the later digit patterning stage. Thus, blastema tissues underwent dynamic changes in their mechanical properties during regeneration, which were different depending on the amputation site.

**Figure 4:**
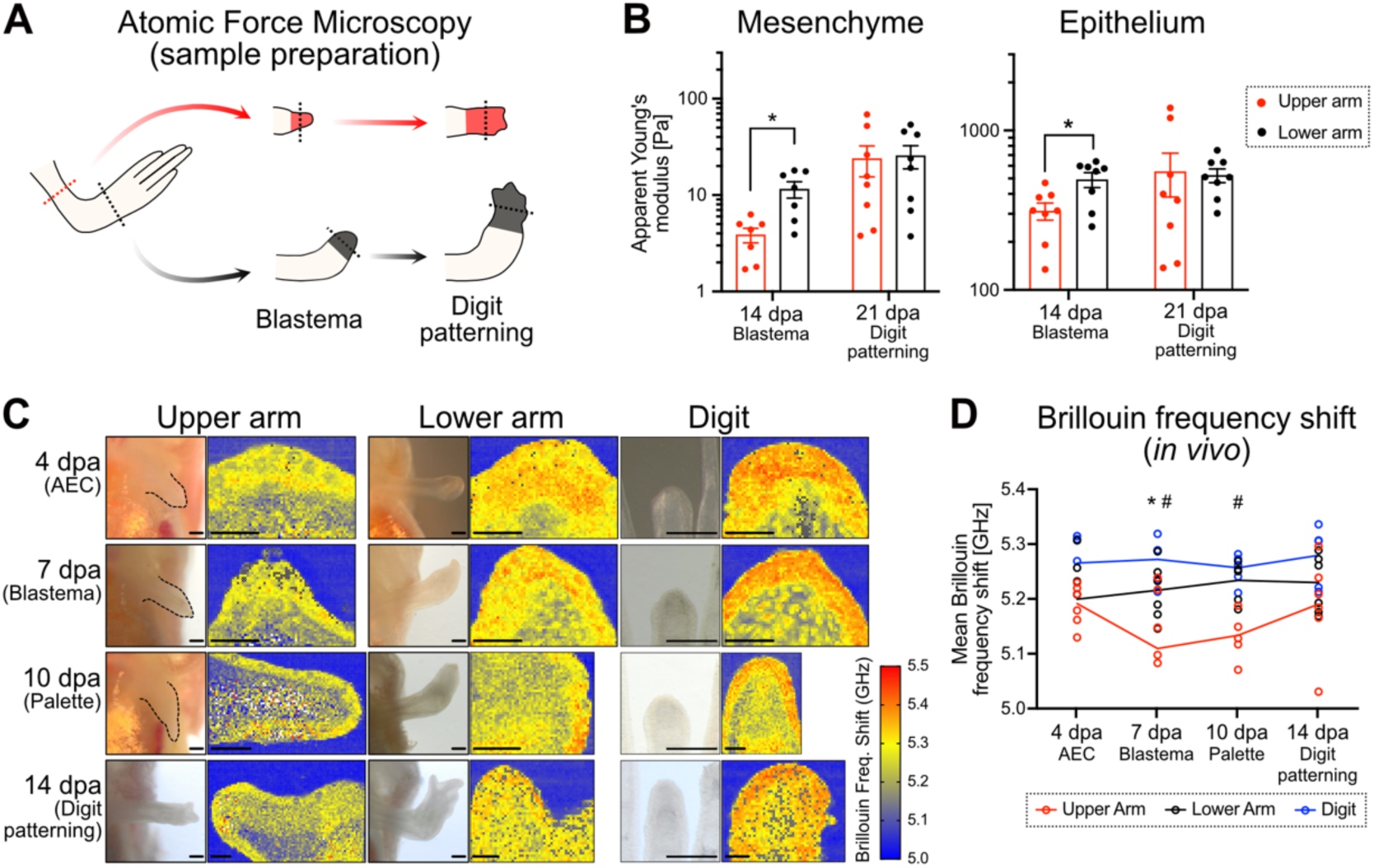
Distal blastemas are stiffer than proximal ones. Limbs were amputated at different levels along the PD axis and allowed to regenerate to probe tissue mechanical properties during regeneration. **A)** Limbs were collected at 2 different regeneration stages and sectioned with a vibratome for AFM measurements. **B)** Apparent Young’s modulus values were measured in the mesenchyme and epithelium by AFM indentation measurements using a cell sized indenter (10 µm radius). Each dot represents the median of at all measurements in one animal (≥ 5 grids, each of 3 x 3 points within 70 µm x 70µm). Columns represent Mean ± SD (n = 8 animals/condition). Mann-Whitney test, * *p* < 0.05. **C)** Brillouin frequency shift (BFS) maps of regenerating axolotl limbs measured *in vivo*. The lower values (blue) correspond to the agarose in which the tissue was embedded, used as external reference. Scale bars: 1 mm (brightfield image) and 100 µm (BFS map). **D)** Mean BFS values during limb regeneration in 2 months-old animals *in vivo.* Each dot represents one animal and lines represent the Mean (n ≥ 5 animals/condition). Two-way ANOVA with Tukey’s multiple comparisons test, \**p* < 0.05 (Upper arm *vs.* Lower arm), ^#^ *p* < 0.05 (Upper arm *vs.* Digit). dpa: days post amputation.

To complement the AFM data, we used Brillouin microscopy, a type of optical elastography that allows the inference of viscoelastic properties of a sample in 3D with high resolution in a contact-free and label-free fashion ^55^, thus enabling mechanical probing *in vivo*. When spatially mapping regenerating limbs (Figure 4C), the Brillouin frequency shift (BFS) changed dynamically throughout the regeneration time course, being higher in distally amputated limbs at the blastema and palette stages (Figure 4D). The same differences were observed in blastemas from larger animals measured *ex vivo* (Figure S5). The BFS is directly related to the longitudinal modulus of the material, which describes its resistance to compression when lateral deformation is constrained and can, therefore, be used as a proxy for tissue stiffness^55^. Thus, BFS measurements reproduce the trend of apparent elastic moduli measured in tissue sections with standard contact-based AFM.

To better align BFS with elastic moduli results, we analyzed regenerating limbs with both Brillouin confocal microscope and AFM by probing the shared surface between adjacent tissue slices (Figure S6A-B). This revealed a relatively high correlation (R^2^: 0.734) between BFS quantifications and apparent Young’s moduli (Figure 4B and S6C), which can be appreciated when plotting these values together (S6D). Therefore, we feel confident that our *in vivo* measurements (Figure 4C-D) truthfully represent tissue stiffness during different phases of limb regeneration.

Altogether, with two different technical approaches, we demonstrate that distal blastemas are stiffer than proximal ones. But these mechanical differences are only detected during the highly proliferative stages of regeneration: blastema and palette.

### Axolotl cell proliferation is mechanically regulated via YAP/TAZ

In order to determine if increased tissue stiffness is causal for decreased proliferation in the blastema, we first evaluated whether any cell-autonomous differences exist between proximal and distal blastema cells. For this, we disaggregated blastemas from *AxFUCCI*-expressing animals and evaluated proliferation *in vitro* in 2D (Figure 5A). The primary cultures indicated no differences in the proportion of proliferating cells between proximally and distally derived blastema cells (Figure 5B). This suggests that differences observed *in vivo* (Figure 2) are highly dependent on their mechanical context (Figures 4, S5, S6). To test the specific impact of the mechanical environment on proliferation in a physiologically relevant 3D context, we developed a new protocol to culture AL1 cells – an axolotl limb cell line – in PEG-heparin-based biohybrid hydrogels ^57^. These gels allow for mechanical modulations that are uncoupled from biomolecular signaling ^57^. We found that increasing hydrogel stiffness – approximately between 200 and 1000 Pa storage moduli ^57^ – reduced the levels of EdU incorporation, indicating a lower proportion of proliferating cells (Figure 5C,D).

**Figure 5.**
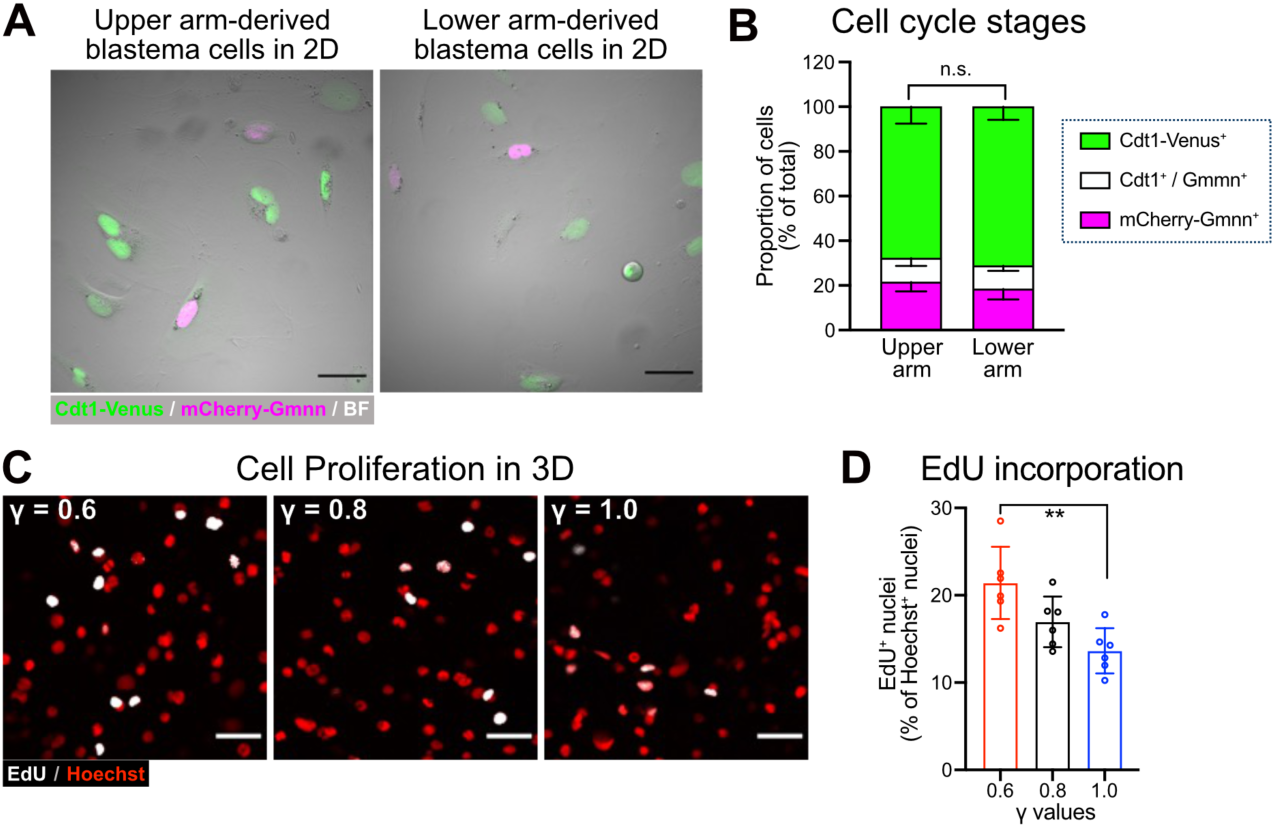
Mechanical dependence for axolotl cell proliferation. **A)** Representative images of cultured blastema cells 48 h after passage derived from *AxFUCCI*-expressing upper and lower arm blastemas. Merged signal of Cdt1-Venus (green) and mCherry-Gmnn (magenta) are shown overlayed on brightfield signal (BF). Scale bar: 100 µm. A) Proportion of cells with detectable signal of Venus (G_0_, G_1_), mCherry (S/G_2_, proliferating cells) or double positive (G_1_/S, G_2_/M, proliferating cells) reveal that blastema cells proliferate irrespective of their PD origin *in vitro.* Mean ± SD is shown (n = 3 wells/condition). Two-way ANOVA, Sidak’s multiple comparisons test, n.s. no statistically significant differences. **C)** AL1 cells were cultured in PEG-based hydrogels with differing degrees of stiffness (higher γ-values correspond to stiffer gels). EdU incorporation was evaluated after 48 h of culture. Representative images of EdU^+^ (white) and nuclei (Hoechst, red) are shown. Scale bar: 100 µm. **D)** EdU^+^ cells with respect to all reveal that axolotl cell proliferation decreases with extracellular stiffness. Mean ± SD (n = 3 gels/condition). Kruskal-Wallis with Dunn’s multiple comparisons test, ** *p* < 0.01.

These results indicate that axolotl cells tune their proliferation in response to the mechanical properties of their niche, with decreased proliferation in stiffer extracellular contexts. To elucidate the causal link between our observations, we studied the activity of YAP/TAZ – known mechanotransducers capable of integrating mechanical cues and transducing them into the coordination of cell proliferation, differentiation, and survival ^58^ – during limb regeneration in the axolotl. For this, we generated a transgenic reporter line consisting of the four times in tandem *GTIIC* enhancer element – which contains consensus binding sites for TEAD transcription factors – upstream of the *EGFP* gene sequence (*4xGTIIC:EGFP*) ^59^.

The *4xGTIIC:EGFP* reporter line reveals YAP/TAZ activity in several tissues throughout the body during early stages of development (stages 46 to 53, based on ^45^) (Figure S7A-K), many of which coincide with its expression in zebrafish ^59^, while others are unique to the axolotl. During early phases of limb development, we detected a clear signal in the interphase between the mesenchyme and the epithelium (stage 46 – 49, Figure S7L-N). At the later digit patterning stages (52-54), the signal from the skin and limb muscles becomes more prominent. (Figure S7O-Q). To validate the *4xGTIIC:EGFP* reporter as a true indicator of YAP/TAZ transcriptional activity in the axolotl, we performed treatments with Verteporfin, a drug that is predicted to restrict YAP/TAZ-TEAD transcriptional activity ^60,61^. We incubated whole embryos in Verteporfin and detected a decrease in EGFP signal intensity in Verteporfin-treated embryos compared to vehicle-treated siblings (Figure S8A-B). correlating well with the expression of YAP/TAZ canonical transcriptional target genes Cysteine-rich angiogenic protein 61(*Cyr61)*, Connective tissue growth factor (*Ctgf)* ^62^, as well as the reporter gene enhanced Green fluorescent protein (*Egfp*) (Figure S8C). These observations suggest that the *4xGTIIC:EGFP* reporter is indeed revealing YAP/TAZ transcriptional activity in the axolotl. Furthermore, we show that local inhibition of Yap activity with Verteporfin in the blastema impairs regeneration (Figure S8D), as expected from evidence indicating that downregulating Yap impairs regeneration as well ^42^.

We used the EGFP signal from the *4xGTIIC:EGFP* reporter as a proxy to evaluate YAP/TAZ transcriptional activity during limb regeneration along the PD axis. For this, we amputated opposing limbs at the upper and lower arm levels and compared intra-individual differences at different regeneration time points (Figure 6A). We detected that the EGFP signal was higher in proximally amputated limbs as compared to distally amputated ones, particularly at the blastema stage (Figure 6B), which suggests that Yap/Taz activity is higher in proximal blastemas. We further looked at regenerating limbs in optical slices of cleared wholemount samples, which corroborated our macroscopic observations (Figure 6C). These observations coincide with the higher *Cyr61* expression in proximally amputated limbs during all evaluated regeneration time points and higher *Ctgf* expression in proximal limb segments in intact limbs, as well as during the blastema stage (Figure 6D). Importantly, these results align with recently reported sequencing data ^63^. As a control for positional gene expression, we used *Prod1* as a proximal marker during early regeneration stages ^21^ and *Hoxa13* as a distal marker, with increasing expression as regeneration progresses ^64^ (Figure 6E). Considering that CYR61 ^65–68^ and CTGF ^69–72^ have been described to promote cell proliferation and survival in several contexts, it is feasible that they are involved in the PD gradient of cell proliferation occurring during early stages of limb regeneration (Figure 1, 2). Hence, we propose a model of growth regulation during limb regeneration in which the gradient in cell proliferation from proximal to distal is a result of differential YAP/TAZ activity, which in turn results from an opposing gradient of tissue stiffness (Figure 6F). However, it remains to be elucidated when and how mechanical differences are established, as well as their connection to PD cell identity.

**Figure 6.**
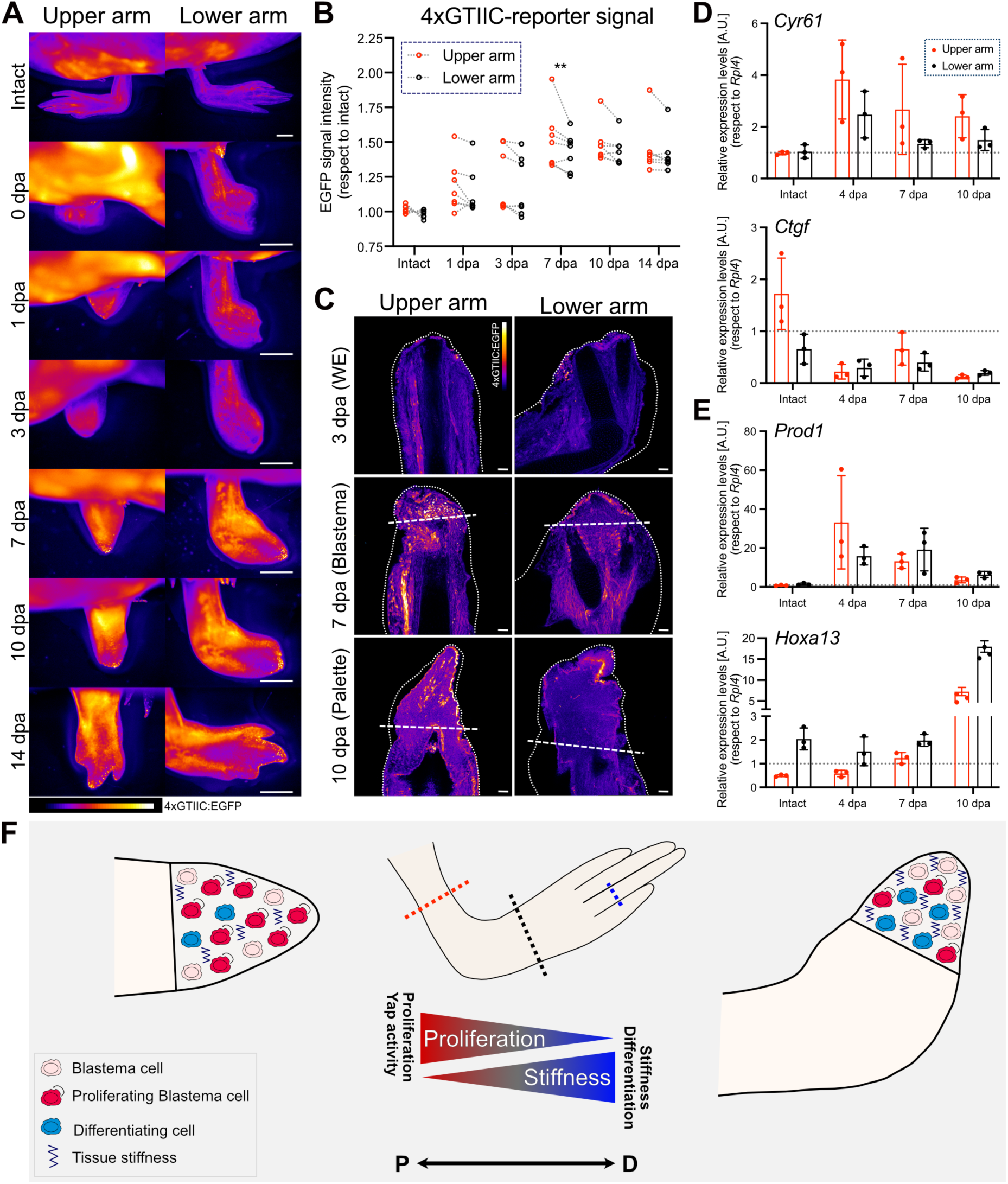
YAP/TAZ activity during regeneration is higher in proximally amputated limbs. Limbs were amputated at the upper and lower arm, after which they were analysed at different timepoints. **A)** Regeneration timecourse from *4xGTIIC:EGFP* animals imaged under a fluorescent stereomicroscope. Scale bar: 500 µm. **B)** Relative EGFP signal from *A*, normalized against the intact condition of each animal. Paired individual comparisons are shown (n = 7 animals). Two-way ANOVA with Sidak’s multiple comparisons test, ** *p* < 0.01. **C)** Wholemount fluorescent imaging of optically cleared limbs at the blastema stage from *4xGTIIC:EGFP* animals. One optical slice is shown. Approximate amputation plane and outline are indicated by a dashed and pointed line, respectively. Scale bar: 100 µm. For *A* and *C*: Both limbs derive from the same animal. Identical brightness/contrast is displayed. Lookup table Fire from Fiji was used to better display intensity differences. **D-E)** Gene expression assessment via qPCR in intact limbs (segments separated at the elbow), at the apical epithelial cap (AEC, 4 dpa), blastema (7 dpa) and palette (10 dpa) stages. Relative expression levels of Cysteine-rich angiogenic inducer 61(*Cyr61*), Connective tissue growth factor (*Ctgf*) (*D*), *Prod1*, and *Hoxa13* (*E*) were normalized against Ribosomal protein L4 (*Rpl4*) levels. Each dot represents one animal. Columns display Mean ± SD (n ≥ 3 animals/condition). dpa: days post-amputation. **F)** Proposed model explaining growth regulation during limb regeneration along the PD axis, in which the gradient in cell proliferation from proximal to distal is a result of a gradient of Yap/Taz activity, which is in turn opposed by a gradient of tissue stiffness and cell differentiation rate. P: proximal, D: distal.

## DISCUSSION

The exact mechanism controlling tissue growth remains a great challenge in developmental biology and regenerative medicine. In this paper, we show that the amputation site directly correlates with the mean cell cycle length and tissue growth rate during axolotl limb regeneration, which is accompanied by important position-dependent mechanical differences. Furthermore, we propose that a YAP-mediated mechanotransduction mechanism is responsible for such observations.

The regulation of growth rate with respect to the extent of tissue removal that we studied here had been previously reported for other body parts and species ^9–13,5–8^, suggesting an ancestral origin of the phenomenon. Whether the same mechanisms are conserved between species remains to be explored. Here, we show that the higher cell proliferation in proximal blastemas is associated with YAP/TAZ activity and lower tissue stiffness, suggesting that growth rate is mechanically regulated during axolotl limb regeneration in a process mediated by YAP/TAZ. These findings agree with studies in the regenerating zebrafish caudal fin showing that there is a gradient of YAP activity in the blastema, from proximal to distal, which correlates with a gradient of cell proliferation ^44^. Furthermore, in *Xenopus* embryos, it was shown that inhibiting TEAD4 decreases growth differences between proximally and distally amputated tails ^13^, although TEAD4 may be required for growth in general, as proliferation was drastically reduced and cell death was triggered regardless of the amputation plane ^13^. Similarly, in the regenerating zebrafish caudal fin, the position-dependent growth rate has been associated with FGF signaling levels in the blastema ^10^, which is preceded by a coupling mechanism between tissue tension and a brief peak in ROS that regulates basal epithelial cell motility during early regeneration stages ^14^. It is possible that such coupling is a result of distinct metabolic states, as it has been reported that the metabolic fluctuations influenced by mechanotransduction pathways may lead to changes in ROS levels and the response to them^73^. In line with this, it has been suggested in the axolotl that ROS is a positive regulator of YAP1 expression and activity during regeneration, as blocking ROS production upon amputation downregulates *Yap1* transcript levels in the limb blastema ^74^ and YAP1 transcriptional targets *Ctgf* and *Areg* in tail blastemas ^75^. This evidence is in agreement with findings in yeast showing that the exogenous application of H_2_O_2_ induces YAP1 nuclear translocation and transcriptional activity ^76^. This data suggests that ROS interacts with mechanotransduction pathways converging in Yap activity regulation; however, whether ROS can directly modulate tissue mechanics is so far unknown.

Decreasing YAP expression or restricting its activity limits growth during regeneration in several species, including the axolotl limb ^42^, zebrafish tail ^44^, and *Xenopus* limb bud ^43^ and tail^13^. Improved regeneration in mice has been achieved by both YAP/TAZ inhibition ^77^, as well as their activation in the skin ^78^ and periodontal tissues ^79^. Overactivation, however, leads to ectopic growth in mouse liver and *Drosophila* eyes ^80^, but increased cell death and reduced size in regenerating zebrafish fins ^44^. This complex plethora of responses indicates that the role of YAP in regeneration is highly context-dependent, which aligns with the fact that the mechanical regulation of YAP activity is far less trivial in 3D environments than shown in 2D *in vitro* ^81,82^. Furthermore, the influence of mechanical forces on cell behavior strongly depends on factors such as biochemical cues and epigenetic state as well ^83,84^.

Here, we report different levels of YAP activity along the PD axis during limb regeneration in the axolotl, which suggests that the regulation of YAP in this context is associated with positional identity. In line with this, it was demonstrated that YAP phosphorylation status, and hence its activity, may be regulated by retinoic acid ^85^, a vitamin A-derived metabolite that regulates PD patterning by specifying proximal cell identities during development and regeneration ^86,87^. ECM mechanical properties may underlie YAP/TAZ distinct activities along the limb’s PD axis as well, as different ECM-related genes are differentially expressed along the axolotl limb PD axis in intact tissues ^20^, during regeneration ^88^, and in response to proximalization induced by retinoic acid treatments ^87^. Interestingly, it is also plausible that YAP plays a role in the determination of positional identity, as YAP inhibition perturbs the expression domains of limb-patterning genes, including a reduction of the distal-associated gene *Hoxa13* and “distalization” of *Hoxa11* in *Xenopus* regenerating limb buds ^43^. It cannot be ruled out, however, that such observations are due to a delay in the regenerative process or decreased proliferation, as the mentioned phenotype is a hallmark of an earlier regenerative stage ^89^.

Mechanical properties drive processes such as cell proliferation, differentiation, and migration in multiple contexts ^29,31–34,38,90–92^. Therefore, it is feasible that the faster differentiation and maturation of cartilage cells observed during the regeneration of distally amputated limbs is a response to the stiffer extracellular environment and decreased YAP activity in distal blastemas as compared to the proximal ones. Supporting this hypothesis is the fact that YAP is a negative regulator of chondrogenesis in mesenchymal stem cells (MSCs) during mouse limb development ^93^.

Tissue mechanical properties may be influenced by active forces driven by cytoskeletal contractile activities, the ECM composition, and cell density ^26,28,30^. Accordingly, tissue stiffness varies greatly throughout the body. For example, fatty tissues and the bone marrow have elastic moduli of tenths of a kilopascal, whereas in calcified bones, they lie within the megapascal range ^94^. Interestingly, we detected relatively low elastic moduli in the blastema mesenchyme (5 - 20 Pa), falling within the range of embryonic tissues ^95^, which – akin to the blastema mesenchyme – are known to present high proliferation rates, as well as being undifferentiated. We detected that tissues in the blastema are significantly softer than during later phases of regeneration, which fits well with former studies indicating that the blastema is softer than mature tissues ^96,97,40,56^. This may be linked to changes in the ECM during blastema formation, as connective tissue cells downregulate ECM-related gene expression, while increasing the expression of matrix metalloproteinases, which breakdown the ECM ^18^. It is possible that this change in extracellular environment and decreased stiffness is linked to the shorter cell cycle detected in regenerating tissues ^48,98,99^. Supporting this hypothesis is the evidence of highly-proliferating cancer cells being 2-3 times softer than their non-malignant counterparts ^100^, and inversely, decreased stem cell proliferation upon niche stiffening during aging ^32,101^. However, increased matrix stiffness has also been associated to the progression of various solid tumors, with matrix stiffness being involved in tumor progression, *i.e.*, increased cell proliferation, invasion, metastasis, angiogenesis, drug resistance, and immune escape (reviewed in ^102^).

Our apparent Young’s moduli values are lower than those reported for blastemas in other laboratories ^96,97^, although in these cases, indentation measurements were performed in bulk using ≥ 300 µm indenter radii and not locally as with AFM. Additional reasons of our value discrepancies may be animal size, age, and the fact that the authors manually removed the epithelium and probed the blastema surface, which means that the boundary between the epithelium and mesenchyme was measured, whereas we did transversal sections with a vibratome and report the values of the blastema center. Furthermore, Kondiboyina and colleagues analyzed formerly frozen samples that were thawed prior to measurements, whereas we measured fresh samples. Until proven otherwise, we cannot rule out that this manipulation may have affected tissue integrity, water content, and consequently, mechanical properties. On the other hand, Calve and Simon performed measurements in a different species, the newt *Notophthalmus viridescens*, which may explain the larger discrepancies between our measurements.

We also report here that the intact limbs from amputated axolotls grow at the same pace as in their intact littermates, which is likely an indication of whole body growth, as limb width is directly proportional to animal body length ^103^. Therefore, animals continue growing regardles of the extent of tissue removal, despite the fact that limb amputations induce a systemic cell cycle entry ^104^. Contrastingly, injured flies and ladybugs delay their development until damaged tissues have regenerated ^46,47^, in a process regulated by Insulin signaling and steroidal hormones ^46,105^. However, to the best of our knowledge, there is no evidence of developmental delays in other species, suggesting that such phenomenon is restricted to insects. The fact that several animals continue growing regardless of the additional energetic requirements that generating missing tissues imply, suggests that important metabolic adaptations take place. In line with this, fluctuating metabolic states associated with cell proliferation, differentiation, and patterning have been documented during regeneration in several species and tissue types ^106–110^, including axolotl limbs ^111^. Considering the reciprocal crosstalk between mechanics and metabolism (reviewed in ^73^), it would be interesting to explore whether distinct metabolic adaptations exist between proximally and distally amputated limbs.

The progress of modern developmental biology has provided great insight into the molecular basis of developmental and regenerative processes. However, only more recently have physical forces and mathematical principles gained comparable attention, likely thanks to the development of technologies allowing their measurement and manipulation ^30^. Examples of *in vivo* studies addressing the impact of external mechanical interventions on regeneration are the mouse digit tip ^38^, cranial bone ^112^ and vascular remodeling ^113^. Regeneration in response to externally applied mechanical cues has been studied *in vitro* as well in *Xenopus* embryonic aggregates ^114^ and mouse intestinal organoids ^115^. Complementarily, physiological mechanical parameters during regeneration have been measured in the zebrafish heart ^116^, spinal cord ^117^ and caudal fin ^14^, as well as axolotl digit ^40^ and limbs ^56,96,97^, among others. Here we show for the first time that tissue stiffness in the blastema is differentially regulated along the limb PD axis, being stiffer in distally amputated limbs. Moreover, we propose that such mechanical differences underlie the regulation of cell proliferation *versus* differentiation decisions during limb regeneration in a process mediated by YAP/TAZ.

Overall, this report, together with a growing body of work, demonstrates how the combination of distinct imaging approaches, mechanical probing, and mathematical modeling can help uncovering the mechanisms behind observations first reported centuries ago. Furthermore, our results provide new insights into the mechanical regulation of tissue growth during regeneration, which may have important implications for the design of biomaterials aimed at being used for regenerative medicine in the future.

## METHODS

### Key resources table

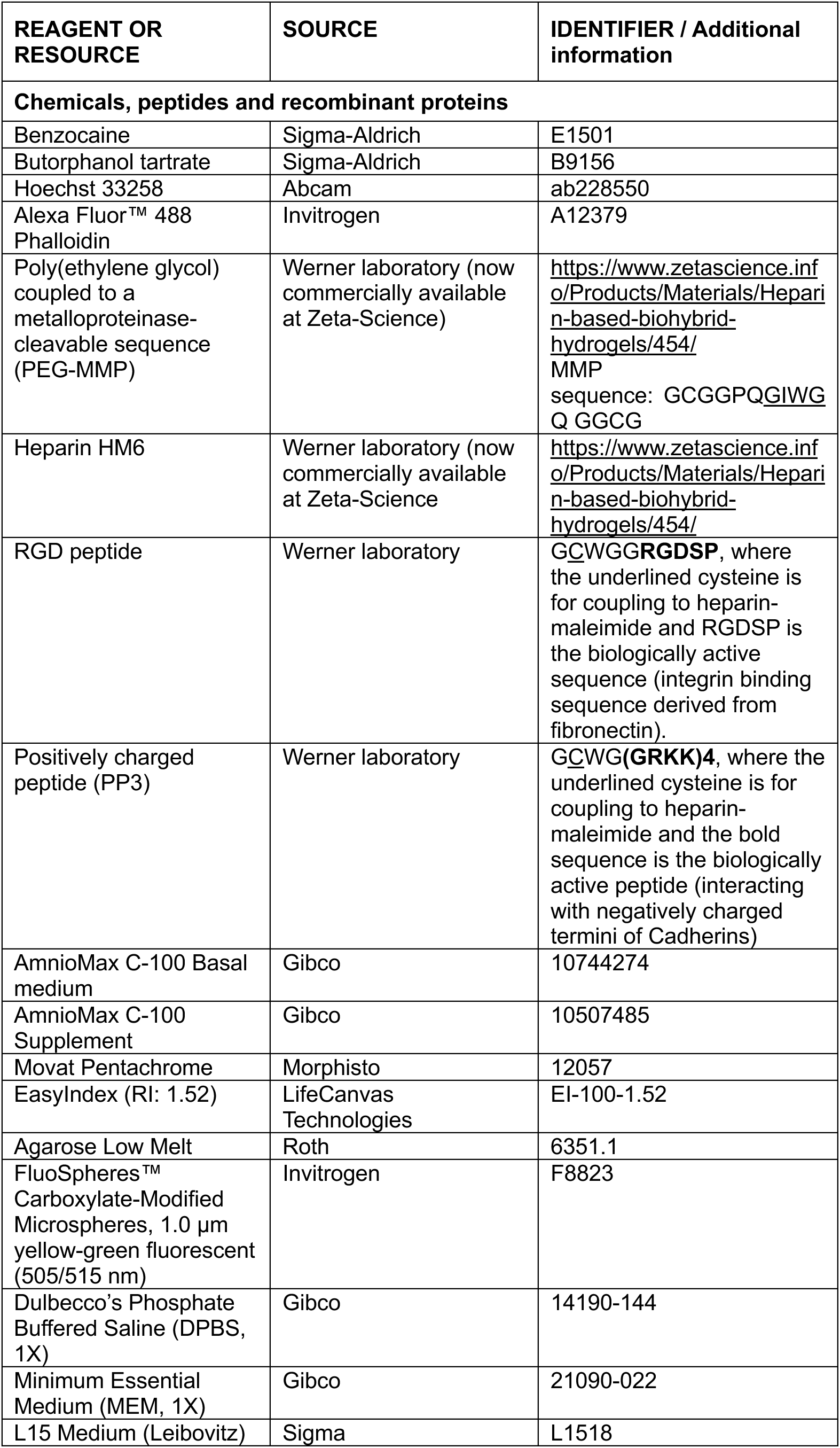
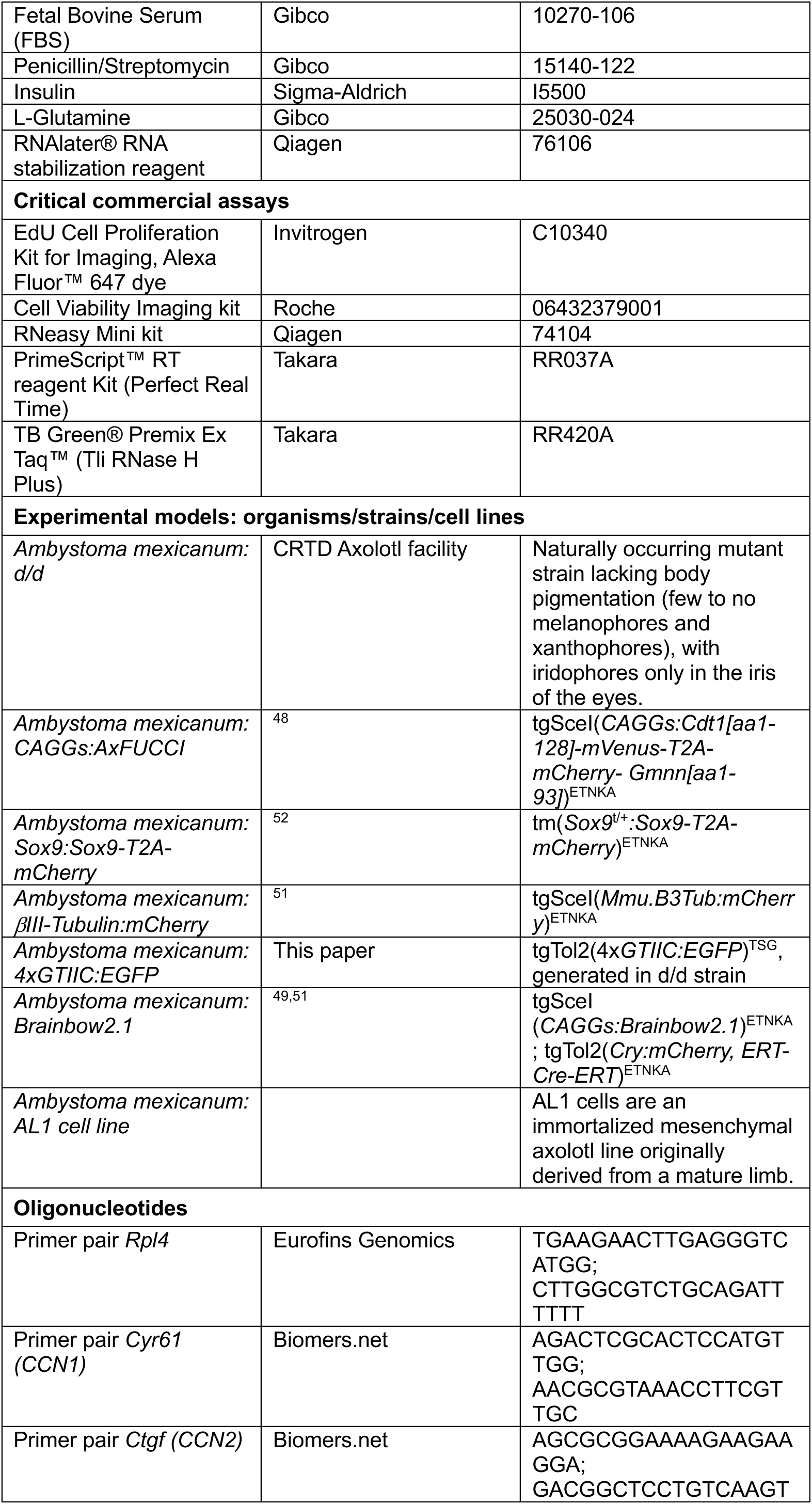
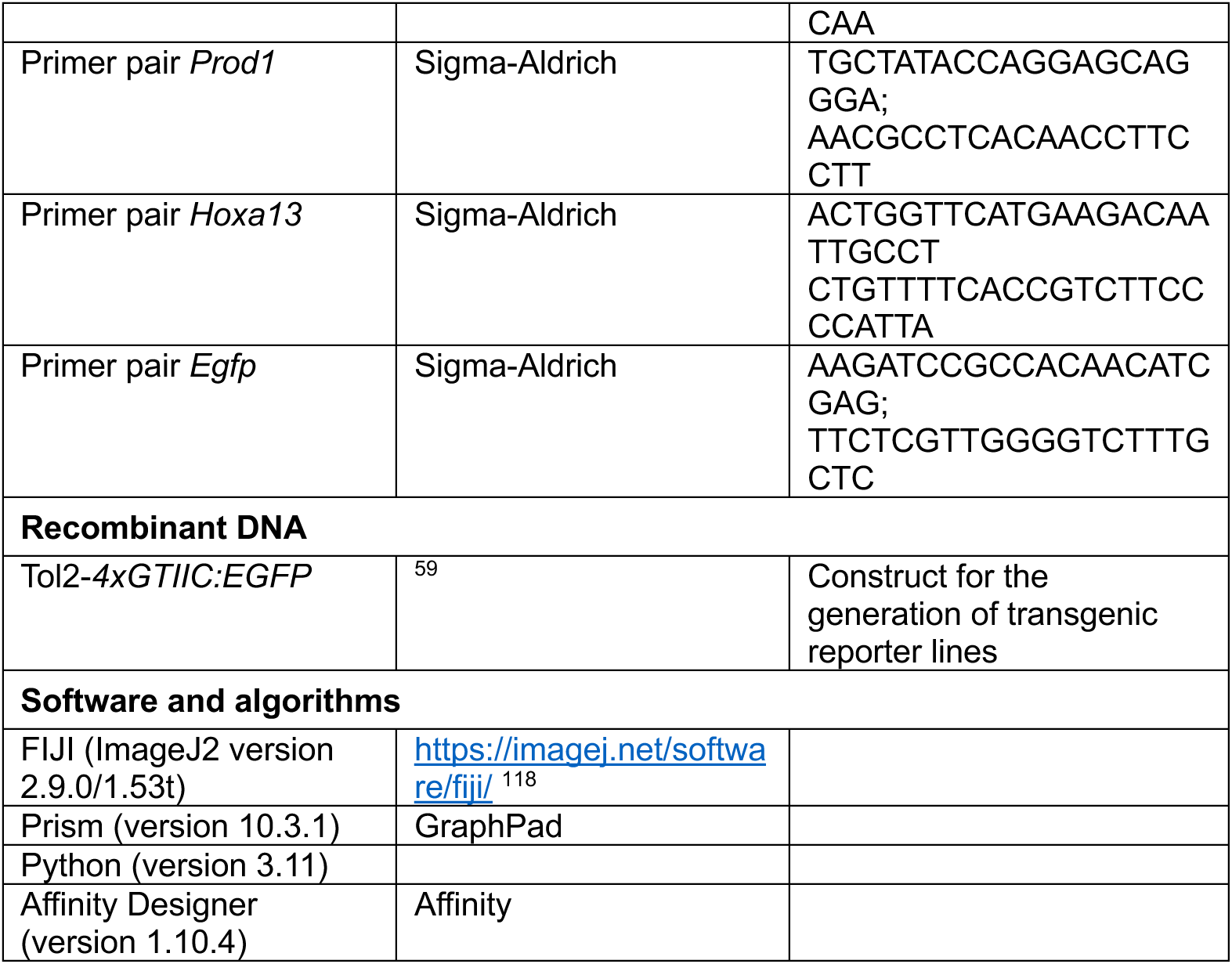

### Resource availability

#### Materials availability

Most of the reagents used are commercially available. All data and transgenic reporter lines generated for this article will be made available upon request.

#### Data and code availability

All custom-made softwares are available on GitHub or Zenodo.

#### Brillouin confocal microscopy

Image acquisition was done with: BrillouinAcquisition v. 0.2.2: C++ program for the acquisition of FOB microscopy datasets (https://github.com/BrillouinMicroscopy/BrillouinAcquisition).

Data analysis was done with: BrillouinEvaluation v. 1.5.3: Matlab program for the evaluation of Brillouin microscopy datasets (https://github.com/BrillouinMicroscopy/BrillouinEvaluation).

Data analysis of discrete areas was done using Impose v. 0.1.2: Graphical user interface for superimposing and quantifying data from different imaging modalities (https://github.com/GuckLab/impose).

#### Mathematical modeling

Repository containing the source code for the analysis and modeling of tissue growth measurements can be found in ^119^ (https://zenodo.org/records/15148889).

### Experimental model and subject details

Axolotls (*Ambystoma mexicanum*) were grown in the axolotl facility of the Center for Regenerative Therapies Dresden (CRTD) of the Technische Universität Dresden (TUD). A full description of the husbandry conditions can be found in ^120^. Briefly, rooms were climatized at 20 - 22 °C with a 12 h / 12 h day/night cycle. All handling and surgical procedures were carried out in accordance with local ethics committee guidelines and were approved by the State Authorities of Saxony, Germany. White (*d/d*) axolotls (*Ambystoma mexicanum*) were used for most experiments, unless otherwise indicated, without any sex-specific bias.

### Generation of *4xGTIIC* transgenic reporter lines

To generate the *4xGTIIC:EGFP* transgenic line, a construct that was kindly donated by Dr. Brian Link ^59^ was used. This is a *Tol2* plasmid containing a synthetic transcriptional enhancer, consisting of 4 multimerized *GTIIC* sequences from the SV40 proximal promoter, which are consensus TEAD binding sites, upstream the minimal promoter of the chicken troponin T (cTNT) gene. This synthetic enhancer drives the expression of EGFP. Fertilized embryos from *d/d* axolotls were injected with the *4xGTIIC:EGFP* vector and *Tol2* mRNA as previously described ^120^. F_0_ animals were selected and grown in the CRTD colony until sexual maturity, after which they were mated with *d/d* axolotls. Experiments were only performed with non-mosaic F_1_ or F_2_ generation.

## Method details

### Animal experimental procedures

A thorough description of animal handling was recently published ^56^. Briefly, prior to all experimental procedures, animals were anesthetized in 0.0075% - 0.01% benzocaine diluted in tap water (higher concentration for larger animals). After collection of tissues or when experiment was finished, animals were euthanized by exposing them to lethal anesthesia (0.1% benzocaine) for at least 20 min. Limb amputations were performed with a sharp scalpel in the center of the calcified area of the humerus, radius/ulna and phalanx for upper arm, lower arm and digit amputations, respectively. After amputations, protruding bones were trimmed with fine dissection scissors. Animals were left in humidified tissue paper soaked with benzocaine-containing water for 15 minutes to allow for blood clotting and wound closure, after which they were returned to their fresh water tank. Painkillers (butorphanol tartrate, 0,5 mg/l) were administered over the first 24 h following surgical procedures.

### Growth measurements and mathematical modelling

#### Limb area measurements

Axolotl limbs were amputated at the upper arm, lower arm and digit levels in 4- and 2-month-old animals (8-9 cm and 4-5 cm long, respectively, measured from snout to tail tip). Weekly images were acquired with an Olympus UC90 stereoscope using CellSense Entry software. Dorsal projection area was measured in Fiji from the joint proximal to the amputation site, of both amputated and contralateral intact limbs, as depicted in Figure 1.

#### Model development

A mathematical model was developed to estimate growth properties in the limb, which required certain assumptions: As cell divisions occur, tissue expands. If cell density can be considered approximately constant, then the number of cells is proportional to the tissue volume. Furthermore, if it is also assumed that tissue depth is constant, then the volume is proportional to the tissue area. The number of divisions at any given time depends on the number of cells present in the tissue at that time. Thus, the rate of change in the number of cells is proportional to the number of cells itself.

If the growth process is unregulated, the tissue could grow to be arbitrarily large. Thus, the regulation of tissue growth was explicitly modelled by requesting that growth rate decreases linearly with the size of the tissue.

The proposed logistic model for tissue area as a function of time is:

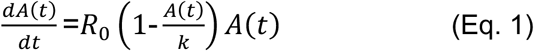

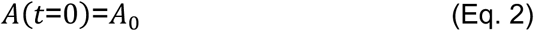

Where *A*_0_ is a constant and represents tissue area at the initial timepoint. The factor *R*_0_ (1-A(t)/k) can be interpretated as the effective tissue growth rate. *R*_0_ is the maximum growth rate possible, *k* is the area at which there is an inflection point and (1-A(t)/k) is the regulation that occurs as the tissue grows.

This system has the following analytical solution:

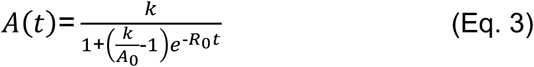

An illustrative example of the area as a function of time obtained by Eq. 3 can be seen in Figure S2E.

#### Fitting the model to experimental data

Data analysis was performed with a custom-made program developed in Python ^119^. We used the function curve_fit from the library SciPy ^121^ to find the optimum parameter values from our model. The mathematical model was fitted to data obtained from each individual animal. To avoid local minima during parameter estimation, a custom multi-start algorithm was developed that randomly chooses the initial guess of the fitting algorithm from parameter values of 4 different orders of magnitude.

#### Model predictions

The cell cycle length θ(*t*) was calculated from the effective tissue growth rate as:

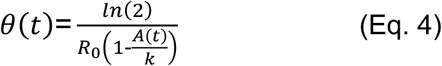

The mean cell cycle length of the entire regeneration process t_f_, was defined as the time integral of θ(*t*) between times 0 and t_f_ divided by t_f_. Importantly, we are assuming that cell cycle length is equivalent in all cell types and we are calculating net values, which means that a combination of cell proliferation and cell death was considered, and that individual contributions of these processes were not calculated.

With the aim to compare the cell cycle length at different phases of the regeneration process, three different time phases were defined: t_1_, t_2_ and t_3_, covering the total duration of the experiment (Figure S2E). Such phases were defined as:

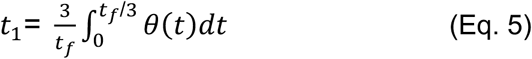

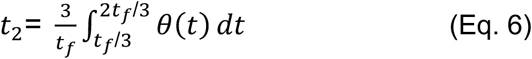

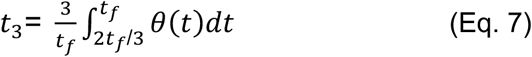

#### Wholemount tissue clearing and imaging

Animals of genotypes *CAGGs:AxFUCCI, Sox9:Sox9-T2A-mCherry, βIII-Tubulin:mCherry*, and *4xGTIIC:EGFP* were used for tissue clearing and wholemount confocal imaging. Briefly, regenerating limbs were collected and fixed overnight in MEMFa (0.1 M MOPS pH 7.4, 2 mM EGTA, 1 mM MgSO_4_ and 3.7% Formaldehyde) at 4°C on a rocking platform. Fixed limbs were delipidated in a solution of 4% SDS and 200 mM Boric acid (pH 8.5) at 37°C, 100 rpm for 30 minutes protected from light. Then, limbs were washed in PBS, followed by nuclei staining with Hoechst (10 µM in PBS) for at least 2 hours at room temperature. After staining, tissues were cleared with an optical clearing solution (Life canvas technology RI = 1.52) at room temperature overnight and stored at 4°C until imaging. Samples were placed on a glass bottom dish (ø: 30 mm) with optical clearing solution and immobilized with a coverslip. Images were acquired with a Zeiss confocal laser scanning microscope LSM780 (Plan apochromat 10x/0.45), with 10 µm between optical planes.

#### EdU injections

To evaluate cell proliferation in large animals, EdU 10 µM mixed with 1% of fast green was intraperitoneally injected (4 µl/g body weight) in anesthetized axolotls. Wounds were allowed to close for 1-5 min ensuring to keep animals moist with wet tissues. Animals were placed back to fresh water and allowed to recover from anesthesia. 4 h after injections, animals were euthanized and limbs were fixed overnight in MEMFa at 4°C on a rocking platform. Fixed limbs were washed thrice with PBS and decalcified in 0.5 M EDTA for 1 week with 2 changes of the solution. After this point, samples were treated as described below.

#### Tissue processing and paraffin sections

Sample embedding, sectioning and staining was performed by the CMCB Histology Facility, Dresden. Briefly, samples were dehydrated in a series of EtOH/dH_2_O until 100% EtOH, and then embedded in paraffin. Longitudinal sections of 5 µm were generated using a microtome.

Movat’s Pentachrome (Morphisto) staining was performed according to the manufacturer’s instructions. Imaging was performed using an Olympus OVK automated slide scanner system (UPLSAPO 10x/0.40).

Proliferating cells were stained with the EdU Cell Proliferation Kit for Imaging (Invitrogen), following manufacturer’s instructions. Cell nuclei were stained with 2 µM Hoechst in PBS. Glass slides were covered with 2 µM Hoechst in a 50% Glycerol solution (in PBS). Slides were imaged in a widefield inverted fluorescent microscope Zeiss Axio Observer.

#### Tamoxifen treatment

To generate clones in the epithelium, a tamoxifen treatment was performed to 2-month-old *Brainbow2.1* transgenic animals. For this, animals were treated with Tamoxifen 1 µM (in holding water) by immersion one day prior to amputation over 30 minutes. After incubation, animals were returned to their tanks containing regular holding water. Regenerating and contralateral limbs were collected at the early blastema and late regeneration stages (6 and 33 dpa), fixed overnight with MEMFa at 4°C on a rocking platform. Fixed limbs were washed thrice with PBS and imaged with a Zeiss confocal laser scanning microscope LSM780 (Plan apochromat 10x/0.45), with 10 µm between optical planes.

#### Cell proliferation assessment

Proliferating cells in *AxFUCCI* limbs imaged in wholemount were manually counted in Fiji. Both the mCherry (proliferating cells) and Hoechst (all nuclei) were counted in at least 3 optical slices in the mid-section of regenerating limbs. The total count was used to determine the proportion of proliferating cells *per* sample. Both the mesenchyme and epithelium were analyzed separately.

To determine EdU incorporation in slices, both EdU and Hoechst channels were automatically counted with the StarDist 2D plugin in Fiji. 8 histological slices *per* sample were quantified, and the calculated proportion of proliferating cells in epithelium and mesenchyme were averaged.

To quantify average clone size in the epithelium of *Brainbow2.1* transgenic animals, all cell clones were manually counted from entire blastemas and digits. To avoid double counting, the Multipoint tool in Fiji was used to label the previously counted cells. All clone sizes were registered manually in a worksheet. Histograms were generated with GraphPad Prism.

#### Whole embryo imaging

Live anesthetized *4xGTIIC:EGFP* embryos were imaged under a Zeis stereoscope using 7.5X - 30X magnification on a petri dish, after which they were immediately placed back in an appropriate container with fresh water. Embryos were imaged at different developmental stages, based on the extended table of stages of axolotl development ^45^.

#### Verteporfin incubations

Drug treatments in freshly hatched *4xGTIIC:EGFP* embryos were performed by incubating embryos in a Verteporfin 20, 50 and 100 µM solution diluted in holding tap water, using the same volume of DMSO as vehicle control. Embryos were incubated for 72 h, after which they were stored in RNA stabilization reagent at 4°C until RNA extraction. Drug treated embryos were imaged as described above immediately before and after drug incubations under a fluorescent stereomicroscope (Olympus UC90 using CellSense Entry software).

Drug treatments in regenerating limbs were achieved by directly injecting the drug (Verteporfin 2 mM) or vehicle (DMSO) solution in the blastema with a microinjector at 5-, 7-, and 9-days post-amputation. Both solutions were diluted in sterile 80% PBS and stained with 1% Fast green. Limbs were imaged weekly over 5 weeks.

#### Gene expression assessment via qPCR

RNA was extracted from whole embryos or regenerating limbs by using the Qiagen RNeasy Mini kit, following manufacturer’s instructions. cDNA was synthesized by using the Takara PrimeScript RT reagent Kit, following manufacturer’s instructions. The resulting cDNA was 5 times diluted prior to gene expression assessment via qPCR, by using the Takara TB Green Premix Ex Taq kit, following manufacturer’s instructions.

### Brillouin measurements

#### Sample preparation

Anesthetized animals or regenerating limbs were mounted on glass bottom dishes and immobilized with low melting point agarose 1% in tap water, and later covered with 0.0075% benzocaine diluted in tap water or 80% PBS, respectively.

#### Brillouin frequency shift measurements

BFS maps were acquired using a custom-built confocal Brillouin microscope previously described ^117^, which has been validated for the measurement of axolotl limbs ^40^. The set-up is based on a two-stage VIPA interferometer. Illumination was achieved by a frequency-modulated diode laser with a wave-length of 780.24 nm. The laser frequency was stabilized to the D2 transition of rubidium 85. The set-up has a CMOS camera (IDS UI-1492LE-M), which allows widefield images to be taken. Image acquisition was done with the custom-made software BrillouinAcquisition (https://github.com/BrillouinMicroscopy/BrillouinAcquisition). Imaging was performed using a Zeiss Plan-Neofluar 20x/0.50 objective, resulting in a spatial optical resolution of 1 µm in the lateral plane and 5 µm in the axial direction. Images were acquired with the Brillouin confocal microscope using a 5 - 10 µm step size and 500 ms acquisition time.

#### Data analysis

Acquired data were analyzed using the custom-made software BrillouinEvaluation (https://github.com/BrillouinMicroscopy/BrillouinEvaluation), and values in each map were exported to a CSV file. Exported BFS maps were graphed as heat maps with GraphPad Prism software.

To calculate average BFS within a defined region of interest, discrete areas from BFS maps were extracted with the custom-made software Impose (https://github.com/GuckLab/impose), values were averaged *per* animal and plotted for multiple comparisons analysis with GraphPad Prism.

### AFM measurements

#### Sample preparation

A detailed description of this protocol is displayed in ^56^, with a few modifications. Briefly, axolotls measuring 15 - 16 cm from snout to tail (7 - 8 months old) were amputated as described above at the upper and lower arm levels and allowed to regenerate until the blastema (14 dpa) and early digit patterning (21 dpa) stages. Regenerating limbs were collected and oriented vertically for embedding in 2% low melting point agarose, diluted in 80% PBS, containing 6 µl/ml of fluorescently labelled beads (FluoSpheres 505/515 nm). After solidification, the agarose blocks containing the tissue were glued (221, Best klebstoffe) on a vibratome stage (Leica VT 1200S), submerged in 80% PBS and sliced (0.9 mm amplitude, speed 0.4 mm/s). The distal 1000-1200 µm of tissue was removed, in order to access a transversal cross section in the half of the regenerating structure (previously measured on stereoscope Olympus UC90 using CellSense Entry software). Measurements were performed on the tissue surface exposed from the agarose block. The remaining block was mounted with surgical histoacryl glue (Braun) onto a 35 mm plastic petri dish (Greiner) and approximately 2 ml of medium (62.5 % L15 medium, 10% heat-inactivated FBS, 1% Penicillin/Streptomycin, 1% Insulin, 1% Glutamine) at room temperature were added to cover the tissue. To distinguish tissue edges from the agarose block, fluorescence imaging was performed to reveal the beads contained within the agarose.

#### Indentation measurements and curve analysis

For AFM indentation tests, a Cellhesion 200 setup equipped with a motor-stage (both from JPK/Bruker) on top of an upright light microscope (Axiozoom, Zeiss) was used. Prior to measurements, the AFM cantilever (arrow T1, Nanoworld with 20 µm diameter polystyrene bead, from Microparticles GmbH) was calibrated using build-in procedures of the microscope software (JPK DP). The petri dish with the tissue block was then inserted into the dish holder (JPK Bruker) and an overview image in brightfield and epifluorescence channels was obtained. Then, a specific region within the epithelium or blastema was selected and an array of force-distance curves was recorded at a comparable indentation depth of approximately 2 - 4 µm, with an approach and retraction speed of 7.5 µm/sec, z length of 50 µm, grid size 70 µm x 70 µm with 3 x 3 points. For every probed region an image was acquired to later relate the obtained stiffness values to a particular region. Thereby fluorescent beads permitted to distinguish well the boundary of epithelium and surrounding agarose.

Force distance curves were analyzed using the Hertz/Sneddon model for a spherical indenter using the JPK/Bruker data processing software and the apparent Young’s modulus was derived. A Poisson ratio of 0.5 was assumed. For each tissue, approximately 5 - 8 different regions were probed.

#### Combined AFM with Brillouin confocal microscopy measurements

In the case of Brillouin frequency shift measurements, the tissue surface that was immediately adjacent to the one used for AFM measurements was used for imaging (Figure S6A). The 600 µm thick tissue slice was embedded on a glass bottomed petri dish with 1% low melting point agarose diluted in 80% PBS, and subsequently covered with modified AL1 cell culture medium. Images with the Brillouin confocal microscope were performed as described above, using a 10 µm step size. After measurements, tissue slices were fixed in MEMFa and stained with Alexa Fluor 488-Phalloidin (1:250) and Hoechst (2 µM) diluted in PBS containing 0.1% Triton X-100. Images were taken with an inverted confocal laser scanning microscope (Zeiss LSM780, Plan apochromat 10x/0.45).

#### Blastema primary cultures

*CAGGs:AxFUCCI* transgenic animals measuring 19 - 21 cm long from snout to tail (1 year old) were amputated at the upper and lower arm levels and allowed to regenerate until the blastema stage (17 dpa). Primary blastema cultures were adapted from a previously published protocol ^49^. Briefly, limbs were sterilized in ethanol 70% and rinsed with 80% PBS, after which they were washed with 80% AmnioMax C-100 Basal medium. Blastemas were collected and moved to a gelatin-coated sterile p35 culture plate containing 200 µl AmnioMax complete medium (171.5 µl of 80% Basal medium supplemented with 28.5 µl of C-100 Supplement), within a larger p100 plate containing sterile water to prevent medium evaporation. 3 days after initiating the culture, additional 100 µl of AmnioMax complete medium were added to the plates. Within the second week after initiating the culture, plates were washed with 80% PBS and filled with fresh 300µl of AmnioMax complete medium and passaged once high confluency was reached in certain regions of the plate to spread them homogeneously. Once 80% confluency was reached, cells were seeded at a density of 1000 cells/well on a glass bottomed 96-well plate coated with gelatin and imaged 48 h after passage under an inverted confocal laser scanning microscope (Zeiss LSM780, Plan apochromat 10x/0.45). Proliferating *AxFUCCI* cells were estimated by fluorescence. For this all mCherry^+^, Venus^+^ and double positive cells were manually counted in 3 wells *per* condition.

### AL1 cell culture in two and three dimensions (hydrogels)

#### AL1 cell culture

Immortalized axolotl AL1 cells were cultured on gelatin-coated flasks using AL1 cell culture medium (62.5 % MEM medium, 10% heat-inactivated FBS, 1% Penicillin/Streptomycin, 1% Insulin, 1% Glutamine) and passaged weekly at a ratio of 1:2. The incubator was kept humidified at 25°C, with 2% CO_2_.

#### Hydrogels

To culture AL1 cells in 3D, ECM-inspired biohybrid hydrogels were used. Specifically, a broadly tunable cell-instructive hydrogel system formed by covalent crosslinking of the glycosaminoglycan heparin and a poly(ethyleneglycol)-(GCGGPQGIWGQGGCG)4-peptide conjugate (PEG-MMP), further equipped with covalently bound adhesive peptide ligands GCWG-RGDSP (RGD) and the positively charged GCW(GRKK)_4_ (PP3) peptide, have been used to perform 3D cultures. The biohybrid hydrogel allows for cell-responsive remodeling due to the matrix-metalloproteinase sensitive GIWG-sequence used for crosslinking and a precise adjustment of hydrogel stiffness. PEG-MMP and Heparin-HM6 synthesis is described in detail ^57^, and can now be commercially obtained (see Key resources table). RGD and PP3 peptides were both generated with solid state peptide synthesis by the Leibniz Institute of Polymer Research Dresden (IPF) as described ^57^.

Each gel was prepared so that 20 µl gels contained 5000 AL1 cells each at a final concentration of Heparin-HM6 1.5 mM, RGD peptide 1.5 mM and PP3 peptide 0.3 mM. The concentration of PEG-MMP varied depending on the γ-value, which corresponds to the molar ratio of the building blocks PEG-MMP and Heparin-HM6, and is thus defining the crosslinking degree and the mechanical properties of the different hydrogels.

To prepare the gels, an initial solution of Heparin-HM6 with RGD and PP3 peptides was prepared fresh and mixed with the cell suspension. Each gel was prepared by individually combining the appropriate volume of a PEG-MMP solution with the previously prepared mix. Cell-containing gels were cultured in AL1 culture media inside a humidified incubator at 25°C with 2% CO_2_.

#### Proliferation and viability assays

To evaluate cell proliferation, AL1 containing gels in were incubated with 10 µM EdU diluted in cell culture media for 1 hour, after which cells were fixed and stained with the EdU Cell Proliferation Kit for Imaging (Invitrogen), following manufacturer’s instructions. Cell nuclei were stained with 2 µM in PBS. Gels were imaged on top of glass bottomed dishes and imaged under an inverted laser scanning confocal microscope (Zeiss LSM780, Plan apochromat 10x/0.45).

To assess cell viability, gels were stained with a Cell Viability Imaging kit (Roche). All reagents were prepared as suggested by the manufacturer. We used a final dilution of Hoechst 33342 (1:2000), Propidium Iodide (1:800) and Calcein-AM (1:1668) in cell culture media and incubated over 1 hour. Immediately after staining, the gels were washed with cell culture media and imaged live under an inverted laser scanning confocal microscope (Zeiss LSM780, Plan apochromat 10x/0.45).

#### Data representation and Statistical analysis

All images were processed using Fiji ^118^. Processing involved selecting regions of interest, merging or splitting channels, measuring signal intensity and improving brightness levels for proper presentation in figures. Maximum intensity projections were done in some confocal images (stated in the respective figure’s legends). When appropriate, stitching of tiles was done directly in the acquisition software ZEN (Zeiss Microscopy, Jena, Germany).

All graphs, Brillouin maps and statistical analyses were performed with GraphPad Prism 10 (GraphPad Software, LLC, San Diego, CA, USA). In all cases, we considered a *p*-value below 0.05 as statistically significant. Specific number of replicates, statistical tests and pos-hoc tests are indicated in the respective figure legends.

All figures were generated with Affinity Designer (Serif Europe, West Bridgford, UK). The area around limb images in Figures 1A, S1A and S8D was cleared with either Fiji or Affinity for a cleaner visualization.

## ACKNOWLEDGMENTS

We thank past and current members of the Sandoval-Guzmán lab for continuous support and companionship during the development of this work. We are particularly grateful to Kerstin Brandt for excellent technical assistance and to Anja Wagner, Beate Gruhl, and Judith Konantz for their valuable dedication to the axolotl care. We would also like to thank Dr. Elly M. Tanaka for providing the *AxFUCCI* transgenic line and Dr. Brian Link for sharing the *4xGTIIC*-based reporter constructs. This work was supported by core facilities of the Technology Platform of the Center for Molecular and Cellular Bioengineering (CMCB) of the TU Dresden, namely the Light Microscopy Facility and the Histology Facility.

## FUNDING

This work was funded by a DFG Research Grant (SA 3349/3-1). S.E-J was supported by the European Union’s Horizon 2020 research and innovation programme (Marie Sklodowska Curie Actions Individual fellowship, grant agreement 101022810) and the Postdoc starter kit from the Graduate Academy of the TUD Dresden University of Technology. A.C. and O.C. were funded by a Biotechnology and Biological Sciences Research Council grant [grant number BB/X014908/1] and O.C. by grant PICT-2019-03828 from the Agencia Nacional de Promoción Científica y Tecnológica of Argentina.

## CONFLICTS OF INTEREST

The authors have no conflicts of interest to declare.

## AUTHOR CONTRIBUTIONS

S.E-J. and T.S-G. conceived the study and acquired funding. S.E-J. designed and performed most experiments, analyzed most data, and wrote the manuscript. T.S-G. advised on the project. N.M.S, A.M.O, R.S. and A.T. assisted with experimental work. K.G, R.S. and A.T. analyzed data. A.C and O.C. developed the mathematical model. U.F. and C.W. provided material and advised on hydrogel use. T.S-G and A.T critically revised the manuscript. All authors proofread the manuscript.

## SUPPLEMENTARY FIGURES AND LEGENDS

**Supplementary Figure S1:**
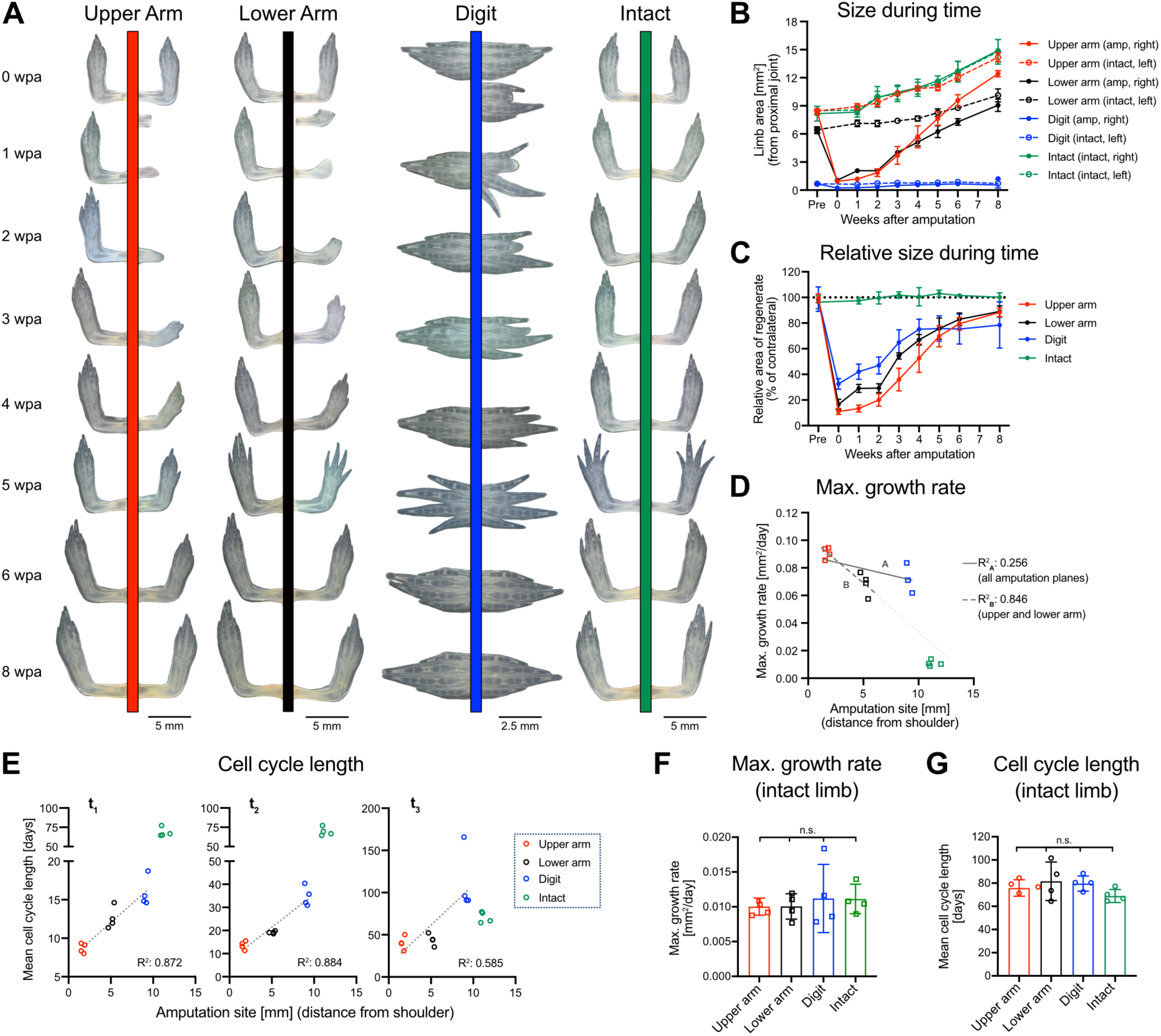
Tissue growth during regeneration is faster after proximal amputations in small juvenile axolotls. 2-month-old animals were amputated at the upper arm, lower arm or digit and regeneration was assessed weekly. One group of siblings were left intact. **A)** Representative images of amputated and intact limbs during 8 weeks. wpa: weeks post-amputation. **B)** Limb area from regenerating and intact limbs (n = 4 animals/condition). **C)** Relative area of regenerating limb with respect to intact contralateral. **D)** Correlation of maximum growth rate with respect to amputation site. **E)** Mean cell cycle duration with respect to amputation site during different phases of regeneration. The entire time course was divided into 3 equivalent segments (t_1_, t_2_ & t_3_). **F-G)** Maximum growth rate (*F*) and mean cell cycle length (*G*) of intact contralateral limbs (measured from shoulder). One-way ANOVA with Kruskal-Wallis multiple comparisons test, n.s. no statistically significant differences. For *B-C, F-G:* Mean ± SD is shown. For *D-G*: Each dot represents one animal. For *D-E*: Intact limbs were graphed according to their length from the shoulder to their longest digit tip. Lines represent linear regression and respective coefficient of determination (R^2^) is indicated (only amputated limbs were included in linear correlations).

**Supplementary Figure S2:**
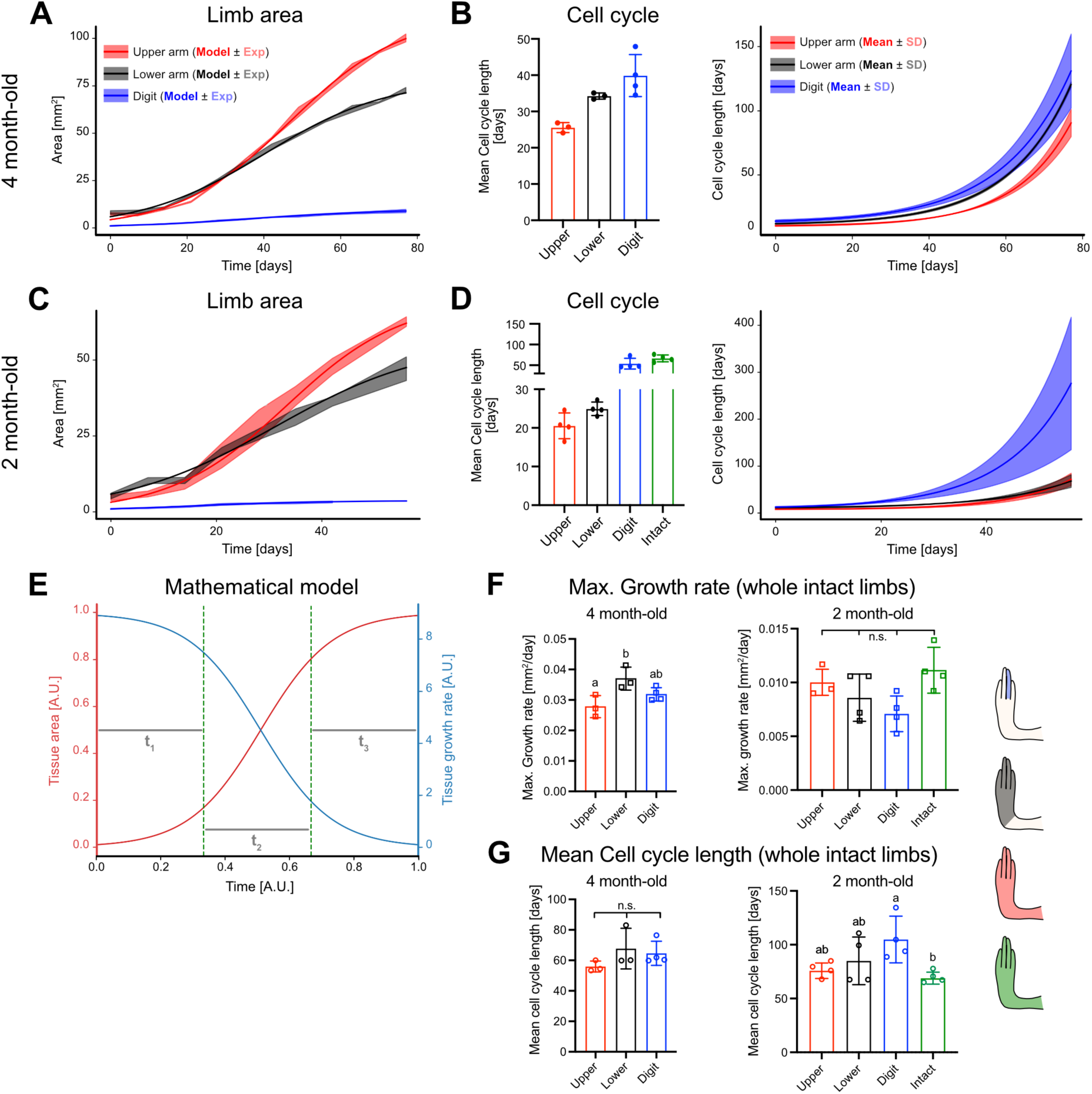
Mathematical model of tissue growth. A logistic regression was used to model limb growth in 2 dimensions during regeneration. **A)** Model adjustment to experimental data for 4-month-old animals. **B)** Average cell cycle length (left) and during the regeneration timecourse (right) from data in *A*. **C)** Model adjustment to experimental data for 2-month-old animals. **D)** Average cell cycle length (left) and during the regeneration timecourse (right) from data in *C*. **E)** Illustrative example of tissue area and growth as a function of time (predicted by Eq. 3), represented by red and blue curves, respectively. At short times, the area increases nearly exponentially because growth rate is not regulated. At larger times, regulation occurs, and smaller tissue growth is seen. Horizontal gray lines mark the time regions t_1_, t_2_ and t_3_. The parameter values to simulate this curve were *k = 1, A_0_ = 0.01* and *R_0_ = 9*. **F-G)** Maximum growth rate (*F*) and mean cell cycle length (*G*) of intact contralateral limbs, measured from the joint most proximal to the amputation site. One-way ANOVA with Kruskal-Wallis multiple comparisons test, n.s. no statistically significant differences. Distinct letters indicate statistically significant differences (* *p* < 0.05). For *B,D,F,G:* Each dot represents one animal. Columns indicate Mean ± SD.

**Supplementary Figure S3:**
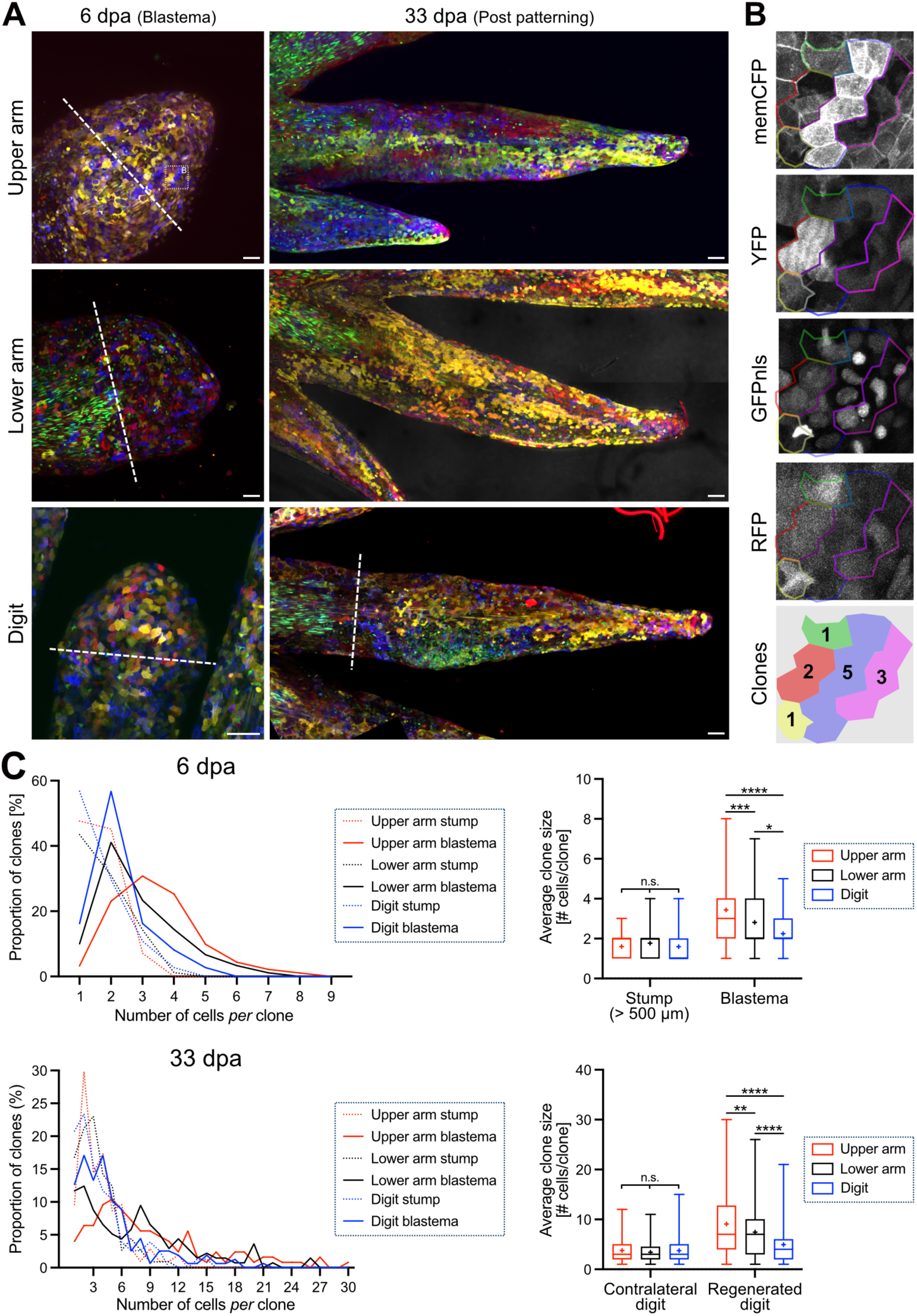
Epithelial cells undergo more rounds of replication after proximal amputations. Limbs were amputated at the upper arm, lower arm and digit levels, after which they were collected at different regeneration timepoints. **A)** Representative images of regenerating limbs of *Brainbow2.1* animals ^49^. The maximum projection of merged signal (membrane CFP, cytoplasmic YFP and RFP, and nuclear GFP) is shown. Scale bar: 100 µm. **B**) Enlarged section from the dotted square in *A* (first quadrant), displaying the different channels for clone size determination. The membrane signal was used to delimitate cell borders. Note that outlines do not coincide perfectly between cortical memCFP signal and the cytoplasmic expression of YFP and RFP. **C)** Proportion of differently sized clones 6 and 33 dpa shown as histogram to display distribution (left) and box and whiskers for statistical analysis (right), with “+” denoting the Mean. Two-way ANOVA, Tukey’s multiple comparisons test, * *p* < 0.05, ** *p* < 0.01, *** *p* < 0.001, **** *p* < 0.0001, n.s. no statistically significant differences (n ≥ 90 clones/animal, N = 1 animal/condition). dpa: days post amputation.

**Supplementary Figure S4:**
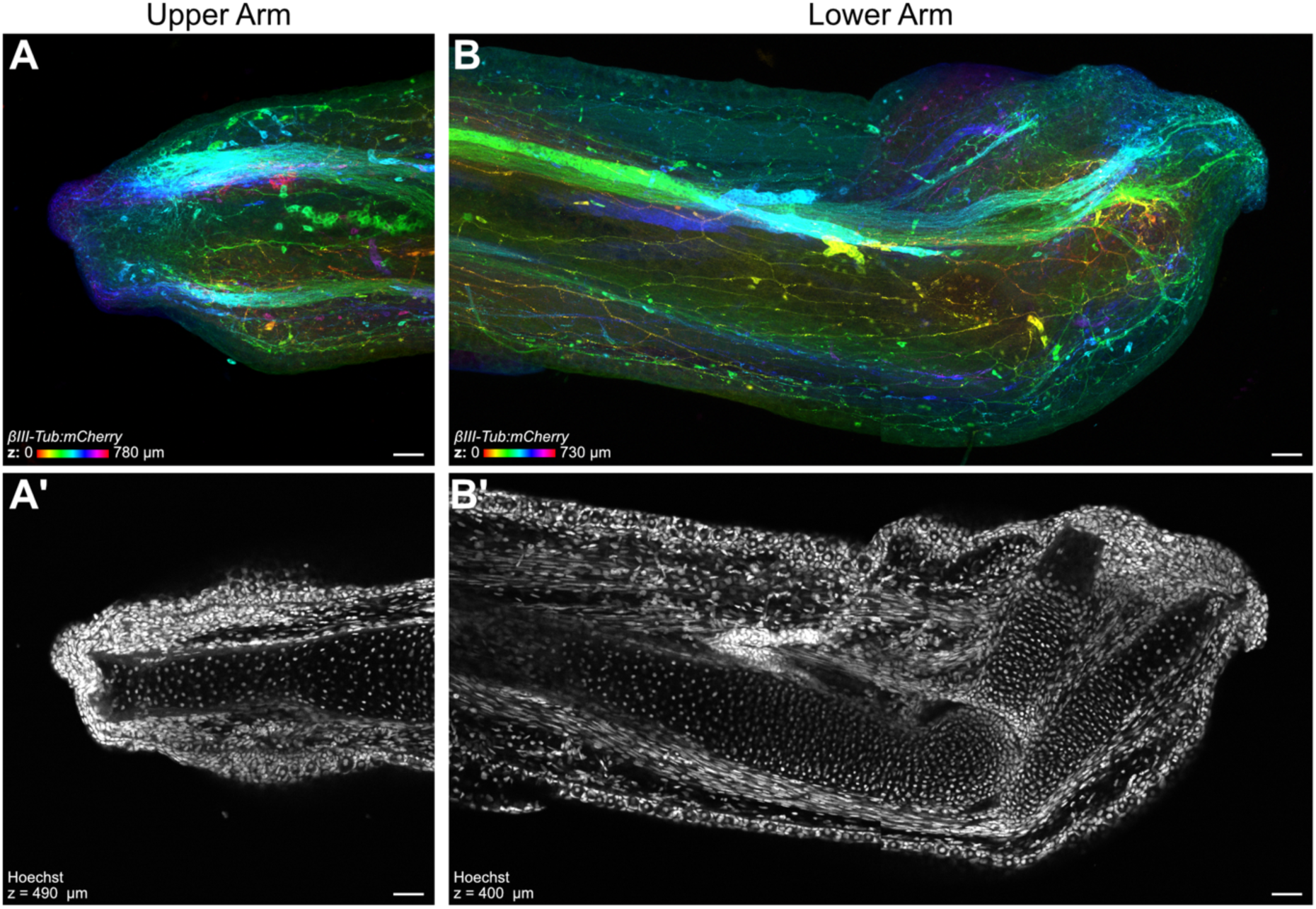
Neural network spreads out more after distal amputations. Axolotls expressing the *bIII-Tubulin:mCherry* reporter of neurons were amputated at the upper and lower arm levels, and allowed to regenerate until the apical epithelial cap stage (4 days post amputation). **A-B)** Maximum intensity projection, color-coded by depth of mCherry signal revealing the neural network distribution. **A’-B’)** Single focal planes displaying nuclei (Hoechst) to reveal bone morphology. Scale bar: 100 µm.

**Supplementary Figure S5:**
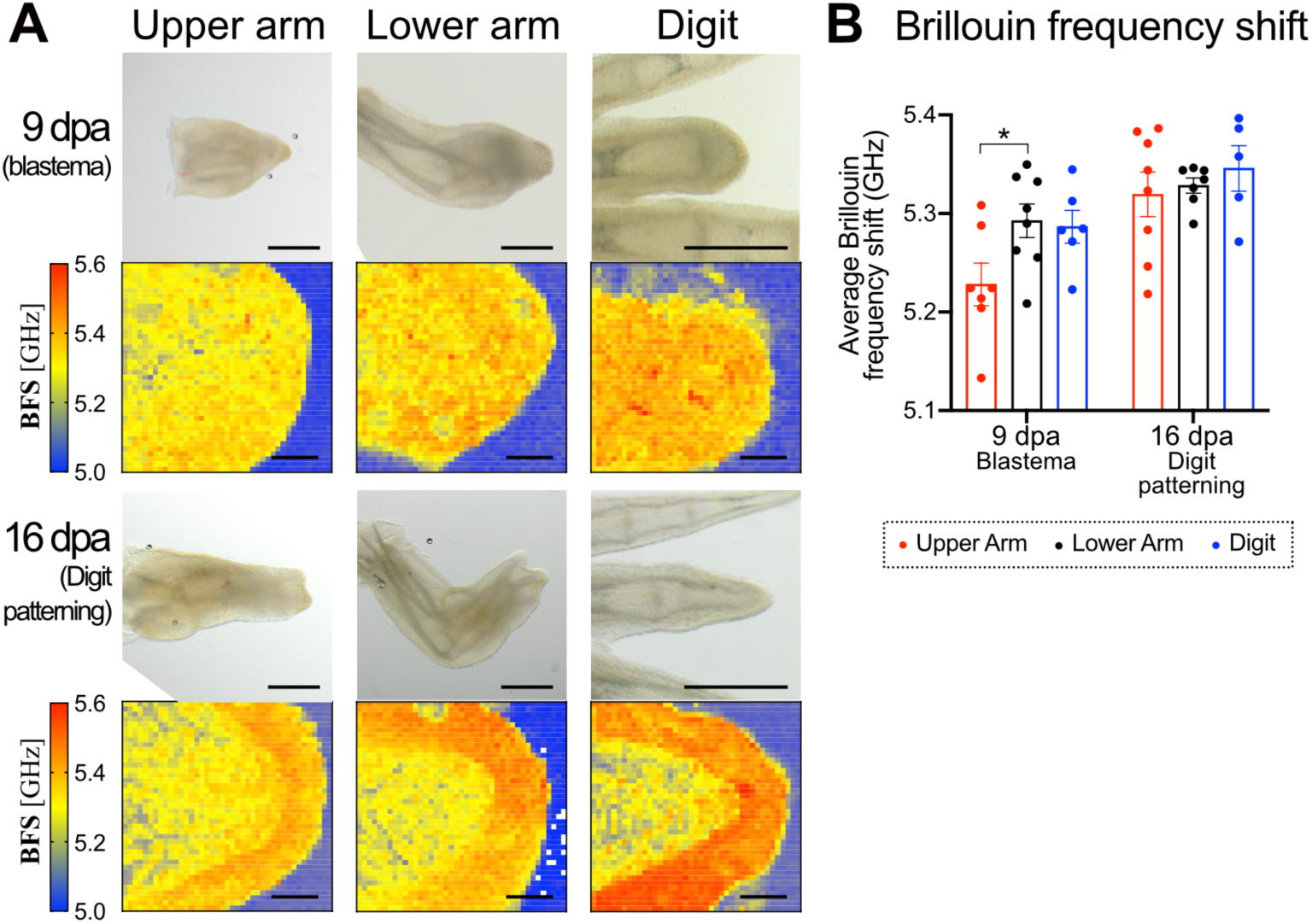
Distal blastemas are stiffer than proximal ones (*ex vivo*). Limbs were amputated at the upper arm, lower arm levels and digit levels allowed to regenerate to probe tissue mechanical properties with Brillouin confocal microscopy. **A)** Brillouin frequency shift (BFS) maps of regenerating axolotl limbs measured *ex vivo* in 4 month-old animals. Scale bars: 1 mm (brightfield image) and 100 µm (BFS map). **B)** Mean BFS values during limb regeneration. Each dot represents one animal and columns represent Mean ± SD (n ≥ 5 animals/condition). Two-way ANOVA with Tukey’s multiple comparisons test, * *p* < 0.05. dpa: days post amputation.

**Supplementary Figure S6:**
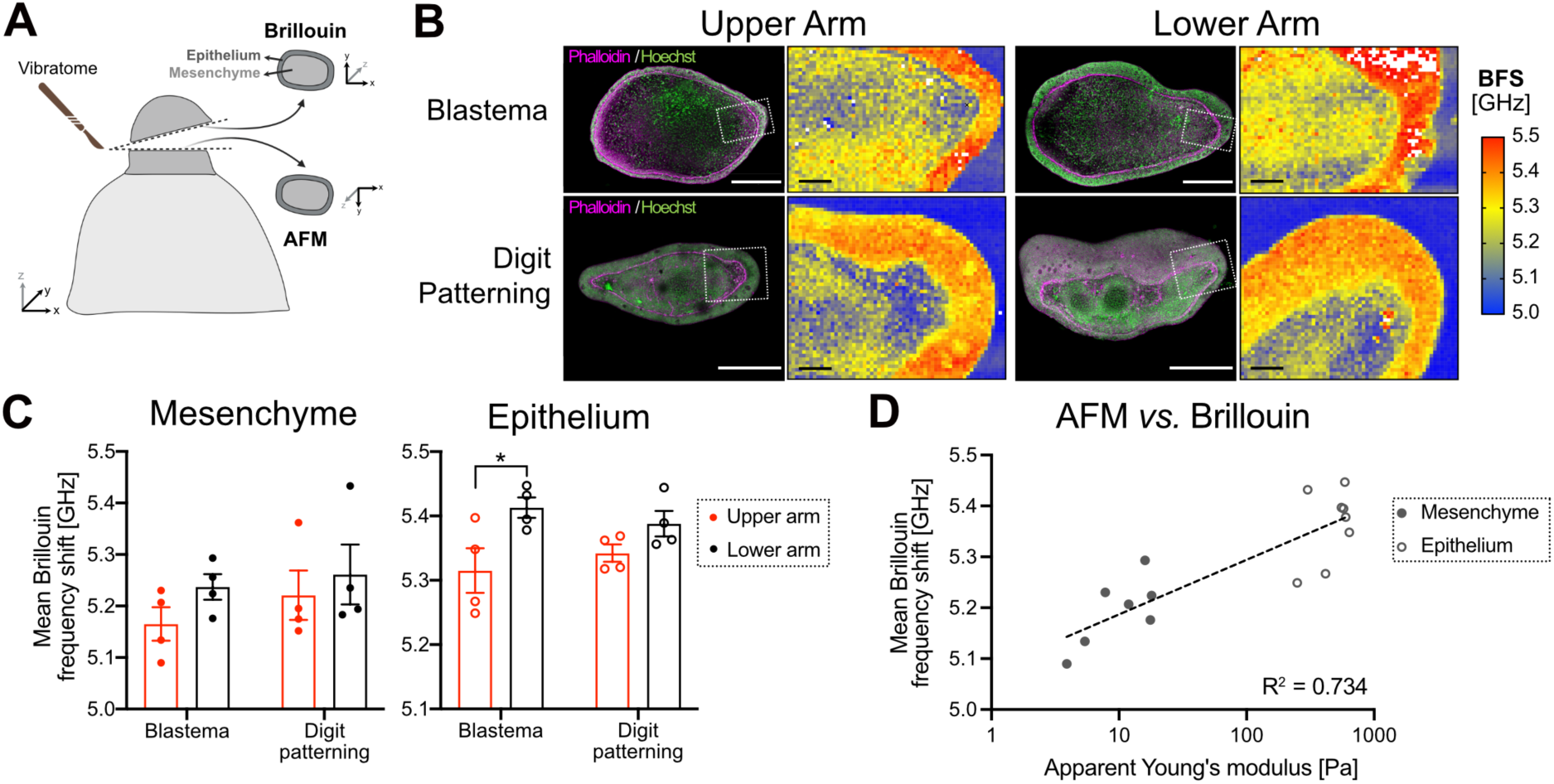
Validation of Brillouin confocal microscopy measurements. Limbs were amputated at the upper arm and lower arm levels and allowed to regenerate until the blastema and digit patterning stages. **A)** Regenerating limbs were sectioned with a vibratome for measurements with the Brillouin confocal microscope and AFM. Adjacent slices were used for cross comparison. **B)** Brillouin frequency shift (BFS) maps of vibratome sections, adjacent to tissue measured with AFM. After measuring with the Brillouin confocal microscope, tissue slices were fixed and stained with Phalloidin and Hoechst to reveal actin filaments and nuclei, respectively. The dashed white square shows the approximate area probed with the Brillouin microscope. Scale bars: 600 µm (immunofluorescence) and 100 µm (BFS map). **C)** Mean BFS values from maps in *B.* Each dot represents one animal and columns represent Mean ± SD (n = 4 animals). Two-way ANOVA with Tukey’s multiple comparisons test, * *p* < 0.05. **D)** Data correlation between AFM and Brillouin confocal microscopy measurements of vibratome slices. A non-linear regression was calculated. The semilog line represents the curve fit and R^2^ the respective coefficient of determination.

**Supplementary Figure S7.**
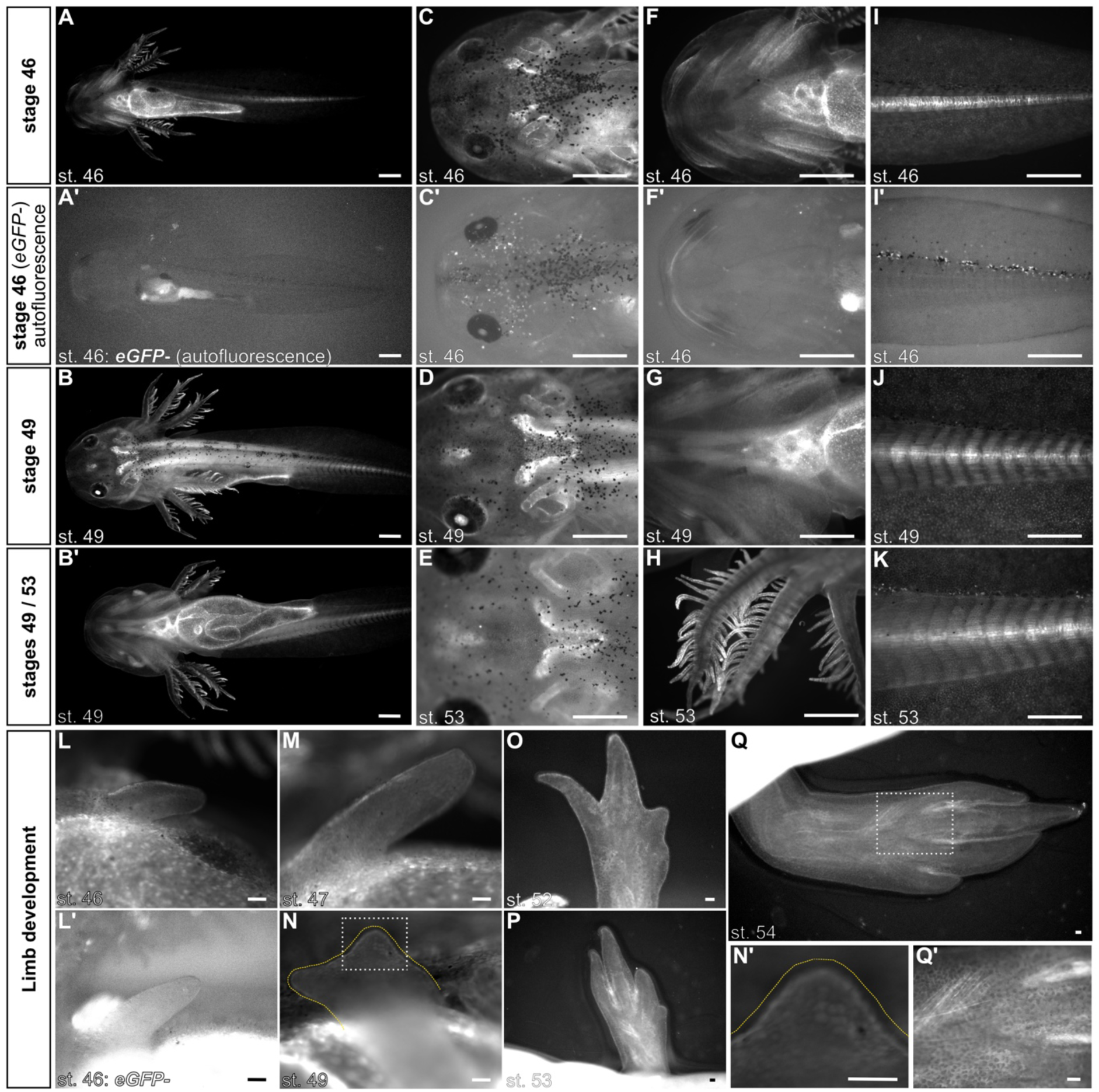
*4xGTIIC*-based reporter expression patterns during development. *4xGTIIC:EGFP* animals were imaged from developmental stages 46 to 54. **A-B)** Whole embryos: Ventral view of stage 46 (*A*), Dorsal (*B*) and Ventral (*B’*) views of stage 49. Signal detected in the skin, dorsal longitudinal muscles (*longissimus dorsi*), and internal thoracic organs, presumably in their epithelium. **C-E)** Dorsal view of the head in stage 46 (*C*), 49 (*D*) and 53 (*E*) embryos. Signal detected in the semicircular canals of the inner ear and what seems to be the *medulla oblongata* or the *nervus laterallis-anterior/nervus octavus,* based on ^122,123^. **F-G)** Ventral view of the head in stage 46 (*F*) and 49 (*G*) embryos, indicating a strong activity in the heart and the lower jaw cartilage. **H)** Gills from stage 53 embryo. **I-K)** Tail from stage 46 to 53 embryos. The notochord is strongly labelled during stage 46 (*I*), and tail segmented muscles (myomeres) show a clear signal from stage 49 onwards (*J-K*). Scale bars from *A-K*: 1 mm. **L-K)** Limbs from stage 46 (*L*), 47 (*M*), 49 (*N*), 52 (*O*), 53 (*P*) and 54 (*Q*) embryos. Signal is detected in the interphase between the mesenchyme and the epithelium at early phases of limb development (*L-N*). At the later digit patterning stages, the signal from the skin and limb muscles becomes more prominent. (*O-Q*). Scale bars from *L-Q*: 100 µm. **N’** and **Q’** show regiones marked in *N* and *Q*. The pointed yellow lines in *N* and *N’* indicate the epithelium, which was drawn using the bright field image. **A’, C’, F’, I’, L)** Non-transgenic animals were imaged under the same conditions as *A, C, F, I* and *L*, respectively, but images are displayed with enhanced contrast to reveal autofluorescence. st: developmental stage, based on table of stages from ^45^.

**Supplementary Figure S8.**
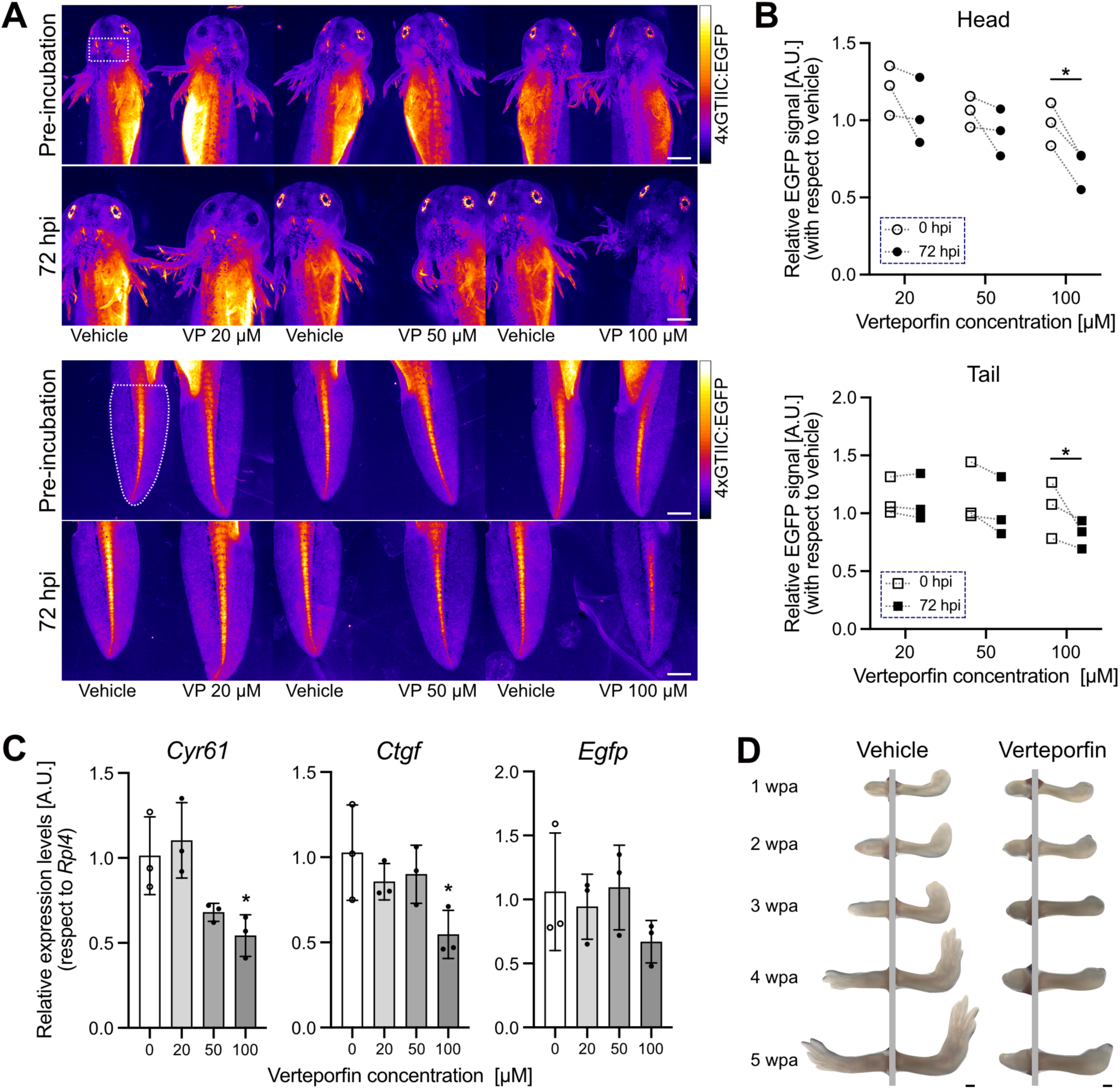
*4xGTIIC*-based reporter validation. **A)** Representative images of embryo heads (dorsal view, above) and tails (below) treated with Verteporfin 20, 50 and 100 µM or control vehicle (DMSO). The same pair of embryos were imaged before and 72 hours post incubation (hpi). Identical brightness/contrast is displayed *per* timepoint. Lookup table Fire from Fiji was used to better display intensity differences. Scale bar: 1 mm. **B)** Relative EGFP signal from area marked by a dashed white outline in *A*. Signal from drug treated animal was normalized against vehicle at 0 and 72 hpi (n = 3 animals). Two-way ANOVA with Sidak’s multiple comparisons test, * *p* < 0.05. **C)** Gene expression assessment via qPCR in vehicle (DMSO)- or Verteporfin-treated embryos. Relative expression levels of Cysteine-rich angiogenic inducer 61(*Cyr61*), Connective tissue growth factor (*Ctgf*) and enhanced Green fluorescent protein (*Egfp*) were normalized against Ribosomal protein L4 (*Rpl4*) levels. Each dot represents one animal. Columns display Mean ± SD (n = 3 animals/condition). One-way ANOVA with Dunner’s multiple comparisons test, * *p* < 0.05. **D)** Representative images of regenerating limbs amputated at the upper arm (left) and lower arm (right) levels. Blastemas were locally injected with Verteporfin 2 mM or control vehicle (DMSO) at 5, 7, and 9 days post-amputation. wpa: weeks post-amputation (n = 6 animals/condition).

## Notes

### Competing Interest Statement

The authors have declared no competing interest.

